# Lipid Flippase Mediated Membrane Asymmetry Governs Extracellular Vesicles Biogenesis and Host Interactions in *Cryptococcus neoformans*

**DOI:** 10.64898/2026.06.12.731820

**Authors:** Siddhi Pawar, Yu Zhang, Christopher Varsanyi, Varsha Gadiyar, Samantha Avina, Raymond Birge, Chaoyang Xue

## Abstract

*Cryptococcus neoformans* is the leading cause of fungal meningitis in immunocompromised patients. Alveolar macrophages are the first line of defense against *Cryptococcus* infection. Our previous study showed that deletion of Cdc50, the regulatory subunit of P4-ATPase (lipid flippase) complex, results in increased phagocytosis and macrophage killing, and avirulence in animal models. However, how fungal flippase dysfunction modulates *Cryptococcus*-macrophage interaction remains unknown. Here we identify Cdc50 as a central determinant of membrane lipid homeostasis, extracellular vesicle (EV) biogenesis and macrophage responses in *C. neoformans*. Our whole cell lipidomic analysis revealed that loss of Cdc50 disrupted membrane lipid homeostasis leading to phospholipid enrichment in *cdc50Δ* mutant, and a reduction in fatty acid production accompanied by pronounced ultrastructural defects in membrane architecture. Loss of Cdc50 also induced a hyper-vesiculating phenotype, with *cdc50Δ* producing significantly more extracellular vesicles (EVs) than wild type H99 cells. Lipidomic profiling of *cdc50Δ* EVs revealed enrichment of phospholipids, including phosphatidylserine (PS), indicating active lipid sorting during vesicle biogenesis. Functional analysis showed that EVs from the wildtype H99 suppress phagocytosis whereas *cdc50Δ* EVs enhance phagocytosis, indicating a differential macrophage priming. Despite increased PS externalization in *cdc50Δ* cells and EVs, macrophage recognition and uptake occur independent of PS-mediated efferocytosis pathways, including PS receptor MertK. Following macrophage uptake, *cdc50Δ* were intrinsically vulnerable to macrophage killing due to rapid phagosome acidification. Together, we demonstrate that Cdc50 dependent lipid homeostasis regulates EV production, lipid composition, membrane architecture and drives the intracellular fate of *C. neoformans*.

**IMPORTANCE:** *Cryptococcus neoformans* is the leading cause of fungal meningitis in immunocompromised individuals. Understanding how this pathogen evades host immune mediated clearance is essential for developing new treatment strategies. Here, we demonstrated that Cdc50, the regulatory subunit of fungal lipid flippase complex, regulates membrane lipid homeostasis that governs extracellular vesicles (EV) biogenesis and macrophage immune responses. Loss of Cdc50 drives global membrane lipid remodeling, hyper-production of phospholipid enriched EVs that enhance macrophage phagocytosis, while the wild-type EV reduce macrophage phagocytosis. Contrary to the prevailing assumption that phosphatidylserine (PS) externalization on the fungal surfaces mimics the mammalian “eat-me signal”, we show fungal PS does not engage canonical PS receptor MertK, revealing a fundamental difference between fungal and mammalian PS biology. Furthermore, *cdc50*Δ cells are unable to resist phagosomal acidification, rendering them susceptible to macrophage killing. These findings establish how phospholipid homeostasis contributes to early host-pathogen interactions and serves as a compelling antifungal target in cryptococcosis.

## Introduction

*Cryptococcus neoformans* is a facultative intracellular pathogen and the leading cause of fungal meningitis in immunocompromised patients, resulting in ∼15-20% HIV-related deaths globally (1). *Cryptococcus* infection is initiated through inhalation of spores or yeasts that are lodged into lung alveoli where they encounter resident alveolar macrophages (2, 3). These early interactions with the alveolar macrophages are pivotal in determining host-pathogen interactions (2, 4). Depending on the host conditions, the interaction can result in pathogen permissive environment, or activation of host protective responses resulting in fungal clearance (5). Although macrophages are well recognized as the critical first line of host defense, the fungal factors that govern macrophage recognition, intracellular fate and immune evasion are still incompletely understood.

Asymmetrical distribution of phospholipids in lipid bilayer membrane is essential for membrane integrity, trafficking, and cell signaling and is maintained by lipid flippase (translocase), lipid floppase (ABC transporters) as well as lipid scramblase (6). Lipid flippase consists of catalytic subunit (P4-type ATPases) and regulatory subunit (Cdc50), which is required for ATP dependent inward translocation of certain phospholipid species (e.g., phosphatidylserine and phosphatidylethanolamine) from exocytoplasmic leaflet to cytoplasmic leaflet (6). *C. neoformans* Cdc50 shares functional and structural properties with both *S. cerevisiae* Cdc50 and Lem3, suggesting evolutionary conservation of flippase regulation across the fungi (7, 8). Lipid flippase functions have been reported in different fungal pathogens. Besides our finding that Cdc50 is critical to antifungal resistance and fungal virulence in *C. neoformans* (9), Hu et al independently demonstrated disruption of lipid asymmetry is accompanied by increased sensitivity to antifungal drugs, drugs associated with phospholipid metabolism, as well as iron chelator, ultimately resulting in severe virulence attenuation in vivo (10). Furthermore, Brown et al demonstrated that Cdc50 influences the activation of Rim101 pathway which helps *C. neoformans* to recognize and respond to extracellular pH and aids in evasion of host immune responses, supporting the notion that plasma membrane asymmetry is involved in response to altered extracellular pH (11). The functional importance of Cdc50 is also highlighted in *Candida glabrata* (12) and *Candida albicans* (13). Loss of Cdc50 in these *Candida* species resulted in hypersensitivity to antifungal drugs, defects in yeast budding, endocytosis, vacuolar functions, increased in cell wall components and activation of cell wall integrity pathway followed by severe virulence attenuation in vivo (12, 13).

Our previous studies demonstrated that phosphatidylserine (PS) is essential for the viability in *C. neoformans* (14) and that the deletion of *CDC50* results in loss of lipid flippase function and PS accumulation on membrane surface (9). As a consequence, the *cdc50*Δ mutant, when co-cultured with macrophage-like cell line J774, showed increased phagocytosis and macrophage-mediated killing, and loss of virulence in murine model of pulmonary cryptococcosis (9). Because in apoptotic mammalian cells PS exposure is an “eat-me” signal recognized by multiple PS receptors on macrophages, including TAM (Tyro3, Axl, MertK) (15) and other PS receptors, such as CD300 families (16–18), it is possible that macrophages may recognize the *cdc50*Δ cells better due to PS accumulation on cell surface through their PS receptors, leading to observed increased phagocytosis. However, whether the fungal PS exposure is recognized by mammalian PS receptors has not been studied. Furthermore, Cdc50 has been implied in pH sensing in *C. neoformans* (11). Because phagosomal acidification is central to macrophage mediated fungal killing, it is also possible that the altered pH adaptation of *cdc50*Δ cells inside macrophages contributes to their susceptibility to increased killing. However, whether defective in acid stress responses contribute to the enhanced clearance of *cdc50*Δ remains to be elucidated.

Emerging evidence on fungal pathogenesis have highlighted the growing importance of extracellular vesicles (EVs) in host-pathogen interactions (19, 20). Cryptococcal EVs are nano-sized, bioactive lipid bound organelles released by the cells in the extracellular milieu that contain lipids, polysaccharides, protein, and nucleic acids (19, 21, 22). These EVs contribute to fungal adaptation to environmental stresses (23), intraspecies, intracellular communications (24) and host interactions (20) (21). These bioactive molecules originate from intracellular compartments and/or plasma membrane (19). The EVs produced by *cdc50*Δ cells may also play a role in promoting macrophage recognition of the mutant cells during yeast-macrophage interaction. Furthermore, functional studies of *S. cerevisiae* lipid flippase Drs2 revealed that the lipid flippase function is required for membrane protein trafficking, and the *drs2*Δ mutant showed less accumulation of ergosterol on plasma membrane and misorting of plasma membrane proteins Pma1 and Can1 (25). Therefore, it is possible that the fungal lipid flippase function may contribute to EV production and function.

In this study, we investigate how lipid flippase, specifically its regulatory subunit Cdc50, mediated membrane asymmetry shapes EV biogenesis and host-pathogen interactions during cryptococcosis. By integrating multidisciplinary approaches, including lipidomic, ultrastructural analysis, EV lipid profiling, macrophage interaction assays and in vivo infection models, we demonstrate that loss of Cdc50 drives global remodeling of membrane lipid composition, pronounced structural defects in the mutant cells followed by increased production of EVs. These EVs differentially modulate macrophage phagocytosis independent of PS mediated efferocytosis pathways. Despite PS accumulation on cell and EV surface, annexin V PS blocking did not alter the phagocytosis. In addition, using a MertK^-/-^ mouse model, we found that the PS receptor MertK does not play a role in the increased phagocytosis in the *cdc50*Δ during its interaction with macrophage *in vitro* and *in vivo* in a murine infection model. Interestingly, *cdc50*Δ is highly sensitive to the pH changes inside the phagolysosome, suggesting its inability to sense and activate Rim101 pathway. Together, our findings establish membrane lipid homeostasis as an upstream regulator of EV biogenesis and determinant of macrophage-mediated killing, elucidating how fungal lipid homeostasis governs early macrophage recognition of a fungal infection.

## Results

### Cdc50 is required for membrane lipid homeostasis

Phospholipid flippase (translocase) is responsible for the inward translocation of certain lipid species, such as PS and PE to help maintain phospholipid asymmetry of the lipid bilayer membrane, a crucial function that underscores membrane stability and supports core cellular processes including vesicular trafficking and cell-surface recognition events (26, 27). In *C. neoformans*, deletion of Cdc50, the regulatory subunit of the fungal flippase, eliminated flippase activity and led to increased phagocytosis and avirulence in vivo (9) (10). However, how the loss of flippase alters membrane architecture and lipid composition in *C. neoformans* has not been studied.

To visualize how loss of Cdc50 affects membrane architecture, we examined the *C. neoformans* cells by transmission electron microscopy (TEM). While wild type H99 and the complement strain of *cdc50*Δ (*cdc50*Δ+CDC50) cells showed a well-defined boundary between cell wall and cell membrane (Fig 1A-B), *cdc50*Δ cells showed loss of clear membrane-cell wall demarcation resulting in membrane invaginations extending into the cytoplasm, appearing as radiopaque membranous structures (Fig 1C-D). Based on our TEM images, only *cdc50*Δ cells showed membrane invagination phenotype. These ultrastructural abnormalities define a mechanically fragile membrane state that aligns with our previously reported sensitivity of *cdc50*Δ mutant cells to SDS and osmotic stressors (9).

**Figure 1.**
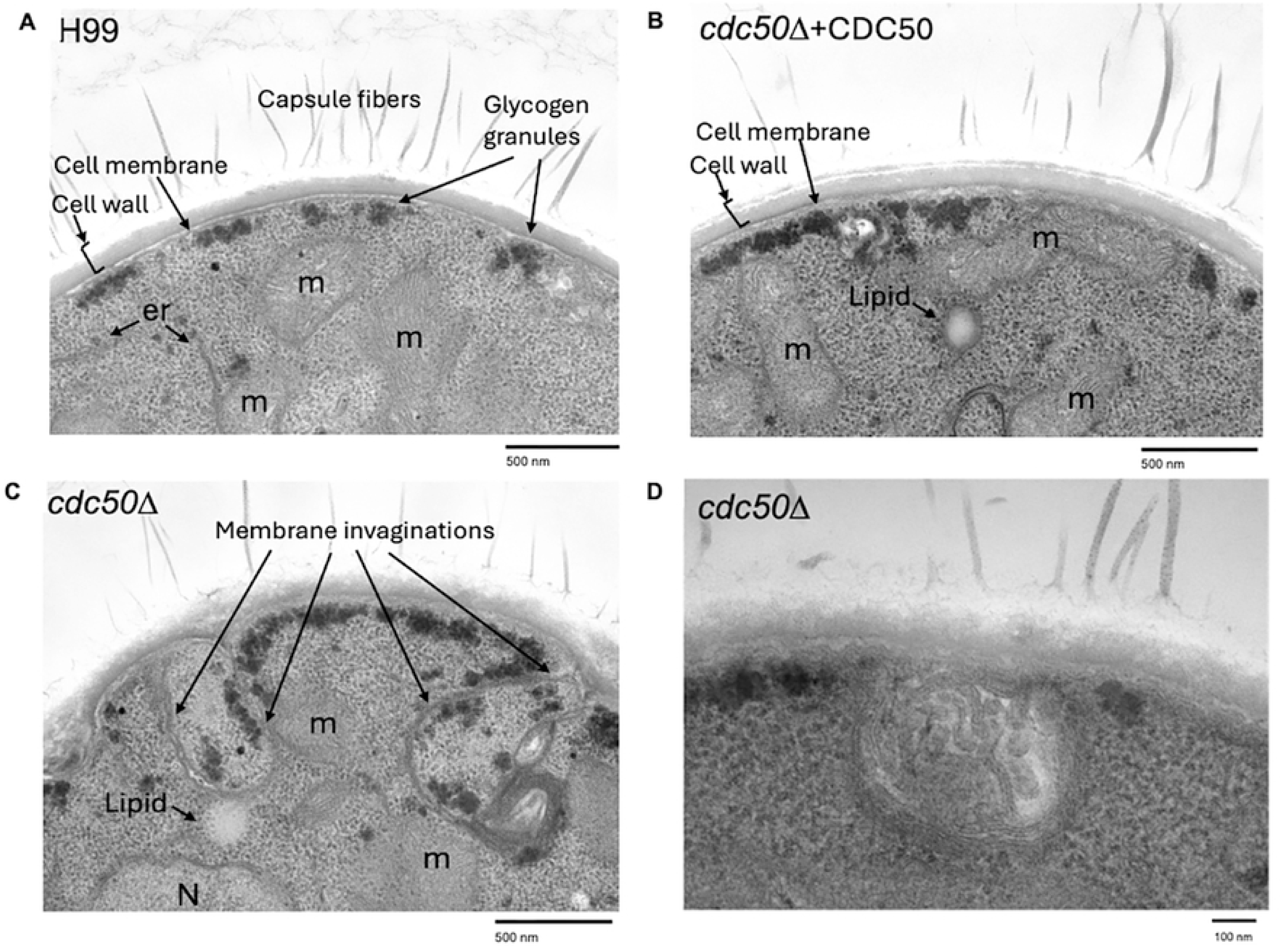
Cdc50 is required for plasma membrane integrity. Yeast cells from wild type H99 (A), *cdc50*Δ (C and D) and its complemented strain (*cdc50*Δ+CDC50) (B) cultured on YPD for overnight were prepared for TEM analysis. Samples were fixed with parafilm blocks before cutting slides. Representative images of each strain were presented and cellular structures were labelled. show a typical morphology of cryptococci of organized membranous compartments and well-defined cell membrane and cell wall decorated with capsule fibers.; (C) Phenotypic traits exclusive to *cdc50*Δ cells include break in plasma membrane, atypical invaginations of plasma membrane followed by electron dense stacked membranes (orange triangle). *cdc50*Δ shows membrane invagination inside the cytoplasm, forming aberrant radiopaque structures (C) & (D)

Next to address how loss of lipid flippase alters membrane lipid composition, we isolated total lipids and performed whole cell lipidomic analysis for H99 and *cdc50*Δ mutant cells using LC-MS/MS-based method. Compared to wild type (WT) H99 cells, the *cdc50*Δ mutant displayed a significant increase in phospholipids (Fig 2A). We observed significant increase in abundance across major phospholipid classes, including phosphatidylethanolamine (PE), phosphatidylserine (PS), phosphatidylinositol (PI), phosphatidylcholine (PC), phosphatidylglycerol (PG), and phosphatidic acid (PA), indicating a global remodeling of phospholipid metabolism in absence of Cdc50 (Fig 2B-G). Furthermore, comparing to H99, we also observed that *cdc50*Δ cells produce much less saturated fatty acid production [FA(13:0), FA(22:0), FA( (19:0), FA( (20:0), FA(15:0), FA(16:0), FA(17:0), FA(14:0) and FA(18:0), etc.,] but more in the monounsaturated fatty acid FA(18:1) and sterols (ST) [ST(24;1;O5) and ST(24:1;O4)]. These observations indicate that the loss of Cdc50 affects fatty acid synthesis essential for membrane stability and lipid homeostasis. The increased phospholipid composition in the *cdc50*Δ mutant cells may be explained by its excessive membrane production and membrane invagination. Collectively, these results demonstrate Cdc50 as a key regulator of plasma membrane lipid homeostasis and that dysregulation of phospholipid distribution is coupled with overt defects in plasma membrane structure.

**Figure 2.**
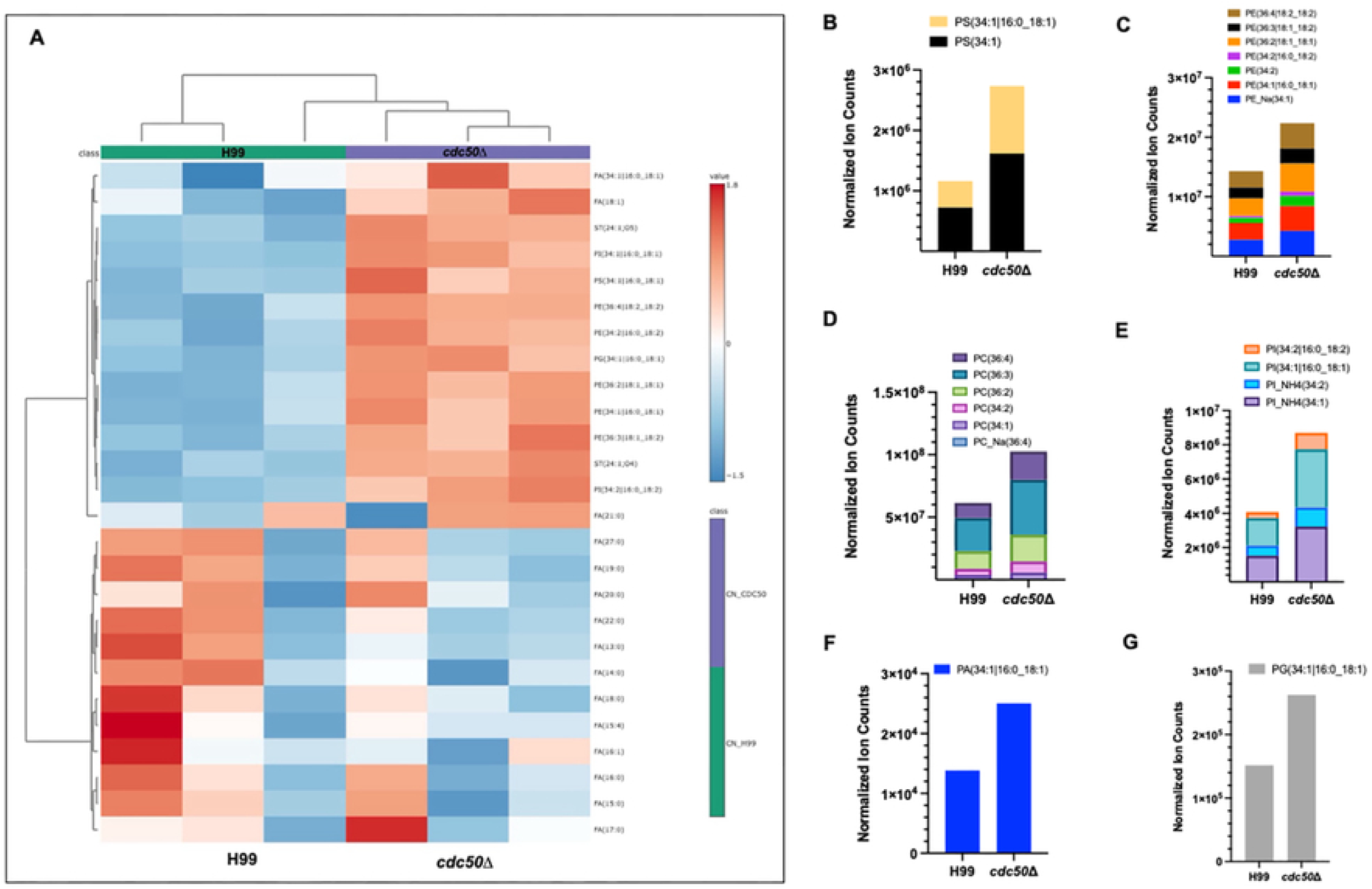
Cdc50 is required for maintaining lipid homeostasis in plasma membrane. (A) Heat map showing differentially expressed lipids in *C. neoformans* H99 and *cdc50*Δ (n=3 for each strain). The major classes of phospholipid species were individually analyzed and compared between wild type and the mutant. (B) Phosphatidylserine (PS); (C) Phosphatidylethanolamine (PE); (D) Phosphatidylcholine (PC); (E)Phosphatidylinositol (PI); (F) Phosphatidic acid (PA) and (G) Phosphatidylglycerol (PG).

### Loss of Cdc50 increases extracellular vesicle production

Our previous studies have shown that deletion of *CDC50* led to increased macrophage recognition and phagocytosis rates (9). Because the *cdc50*Δ mutant has PS accumulation on membrane surface and PS surface exposure has been defined as the “eat-me” signal in apoptotic cells to induce phagocytosis and macrophage-mediated clearance, we hypothesize that the PS exposure in *C. neoformans* cells may also enhance macrophage recognition, leading to increased phagocytosis (9). Because capsule and cell wall may act as a barrier preventing direct PS-PS receptor interaction, we further hypothesize that the *cdc50*Δ cells might secrete PS-rich extracellular vesicles (EVs) to attract macrophages, leading to increased phagocytosis of *cdc50*Δ cells. To test this hypothesis, we isolated EVs from solid cultures of wild-type H99 and the *cdc50*Δ mutant (Fig 3A) and quantified vesicle release by nanoparticle tracking analysis (NTA) as described previously by Reis et al. Our results showed that from the same amount of H99 and *cdc50*Δ yeast cells, *cdc50*Δ cells produced 2.5x 10^11^ EVs/ml while the WT H99 cells produced 0.5x 10^11^ EVs/ml, indicating *cdc50*Δ mutant released five times more EV particles than H99 (Fig 3B). However, the size distribution of EVs produced by the two strains were comparable at 150 nm in average (Fig 3C), indicating that loss of Cdc50 increases vesicle production without any alterations to the vesicle size.

**Figure 3.**
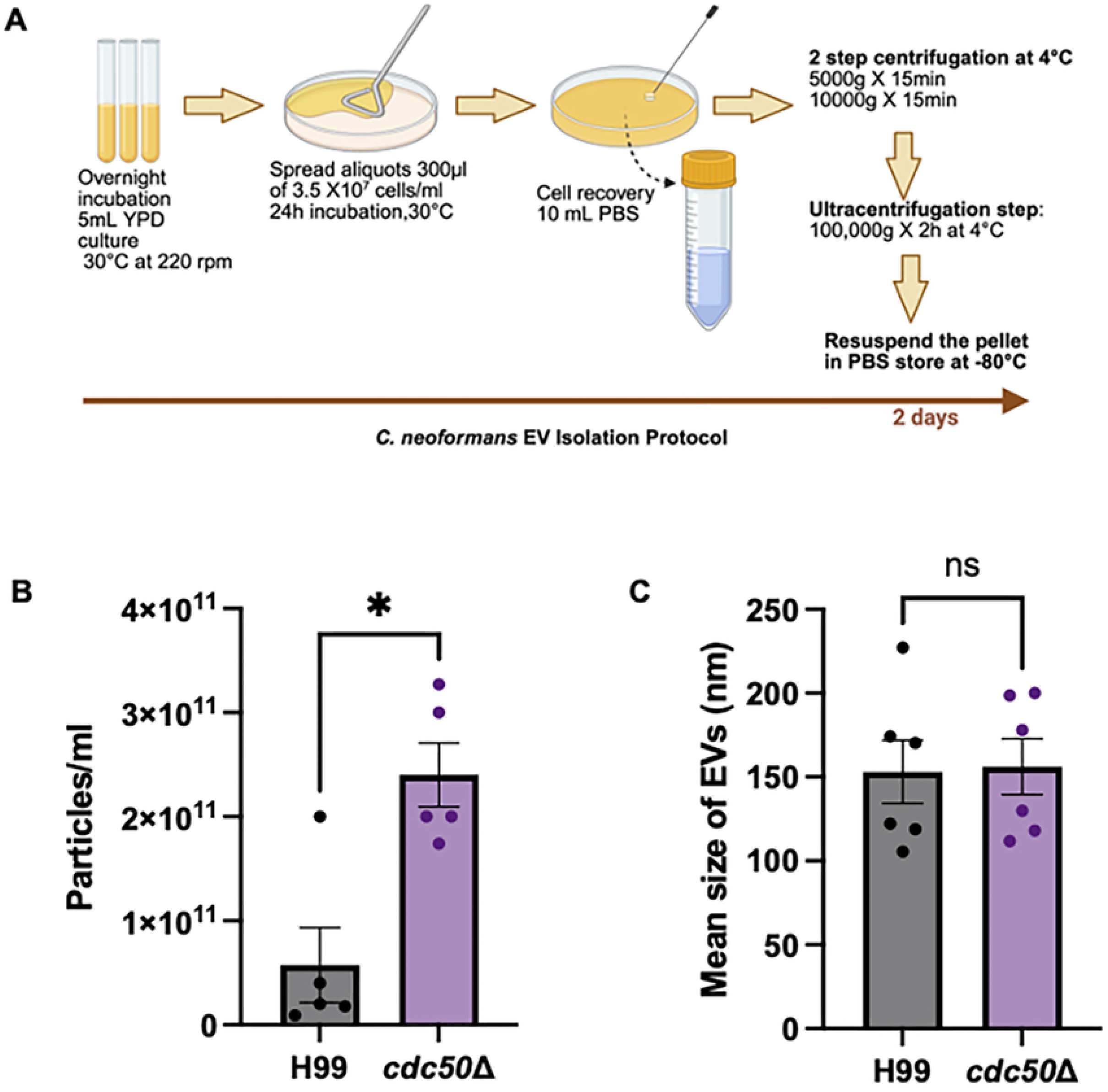
Loss of Cdc50 increases extracellular vesicle (EV) production. (A) Schematic illustration of isolating fungal extracellular vesicles (EVs) from solid cultures. (B and C) Nanoparticle tracking analysis (NTA) to measure (B) concentration and (C) size of the vesicles. Each dot represents an individual sample and bars indicate mean±SEM. Statistical significance between two independent groups for (B) was determined using a Mann-Whitney test (p<0.05) and for (C) was determined using Welch t test ns=no significant difference observed

To further examine EV morphology and membrane structure, we analyzed EVs using TEM. Negative-stain TEM showed that EVs isolated from both strains were membrane-bound vesicles (Fig 4A-B). Ultrathin-resin section TEM further confirmed the presence of membrane enclosed vesicles in both strains. Moreover, H99 EVs appeared predominantly spherical while the *cdc50*Δ mutant appeared more frequently a cup-shaped morphology. However, the cup shaped morphology might be an artefact produced during the EV processing for TEM (22). To visualize the EVs under near-native conditions, we also performed cryo-electron microscopy (Cryo-EM). Cryo-EM confirmed the presence of lipid bilayers in EVs and revealed fibrillar decorations associated with EVs in both H99 and *cdc50*Δ mutant cells (Fig 4C). Together, these results demonstrate that loss of Cdc50 leads to increased EV production while preserving key ultrastructural features of cryptococcal EVs, suggesting that the *cdc50*Δ mutant exhibits a hyper-vesiculating phenotype rather than a catastrophic defect in EV architecture.

**Figure 4.**
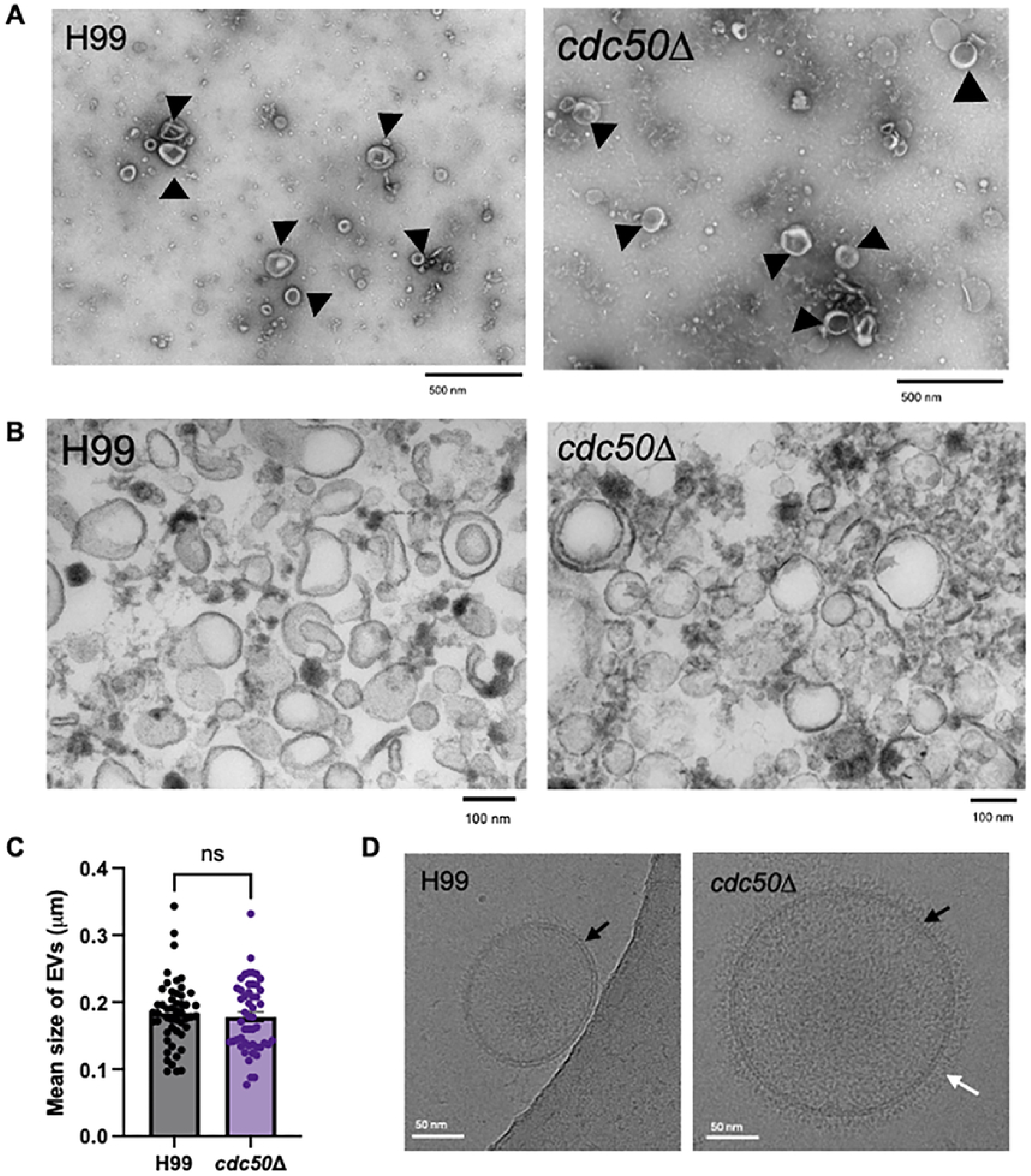
Loss of Cdc50 does not affect the structural architecture of extracellular vesicles. (A) Negatively stained EVs with 1% uranyl acetate show presence of lipid bilayer (black arrow) for both WT H99 and *cdc50*Δ. (B) Ultra-thin resin sections of EVs highlight a cup shaped morphology of *cdc50*Δ with a well-defined lipid bilayer compared to H99 EVs which appear more spherical. (C) Each dot represents a single extracellular vesicle (EV), and the bar plots indicate mean±SEM. No significant difference was observed between H99 and *cdc50*Δ EVs, as determined by Mann-Whitney test. (D) Cryo-EM imaging showing WT H99 and *cdc50*Δ EVs delimitated by lipid bilayer (black arrow) followed by fibrillar decorations (white arrows).

### Loss of Cdc50 alters EV lipid composition and enriches EVs in phospholipids

EVs can originate from endosomal compartments or bud directly from plasma membrane and therefore may reflect the lipid composition of the membrane of origin (28). Our previous study showed increased PS exposure on the *cdc50*Δ cell surface (9), we hypothesized that the EVs released by the *cdc50*Δ cell might also alter PS expression on the EVs.

To test this hypothesis, we isolated EVs from solid agar cultures of H99 and *cdc50*Δ and stained them with lipophilic membrane dye DiL to identify the presence of EVs and annexin V to detect the surface exposed PS (Fig 5A). EVs were then subjected to flow sorting. Our results showed that EVs produced by *cdc50*Δ cells expressed a significantly higher proportion (18.3%) of Annexin V positive vesicles. Specifically, PS-positive EVs in H99 increased from approximately EVs from 5% to 25% in *cdc50*Δ, indicating a five-fold increased PS exposure on the vesicle surface (Fig 5B).

**Figure 5.**
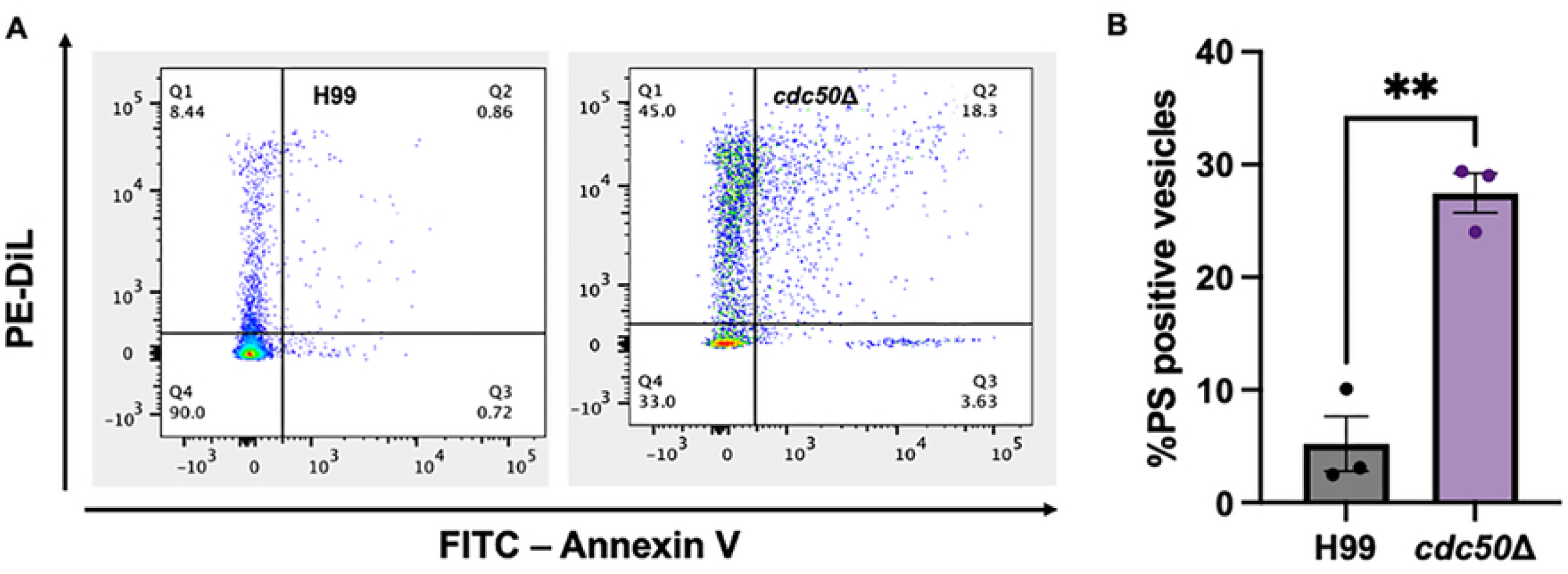
*cdc50*Δ EVs show increased phosphatidylserine on its surface. (A) Gating strategy for the EVs stained with DiL and Annexin V-FITC dye. (B) FACS analysis of WT H99 and mutant *cdc50*Δ show greater percentage of phosphatidylserine on the surface of *cdc50*Δ as compared to H99. Bars represent mean±SEM. ** Statistical significance was determined using unpaired t test p<0.05.

To further determine whether increased PS surface exposure reflected a broader alteration in EV lipid composition, we next performed lipidomic analysis of purified EVs. Total lipids from EVs produced by H99 and *cdc50*Δ cultures were extracted and analyzed using LC-MS/MS-based lipid-omics method. EVs produced by *cdc50Δ* cells showed higher abundance of phospholipids but reduced short chain fatty acid composition than H99 EVs, indicating loss of Cdc50 substantially alters EV lipid composition (Fig 6A). Phospholipid class profiling of *cdc50*Δ EVs showed an enrichment of major phospholipid profiles, PS, PE, PC, PI, PA, and PG, compared to H99 EVs (Fig 5B-G). Notably, PA(34:1|16:0_18:1) (Fig 6F) and PS(34:1|16:0_18:1) (Fig 6B) were enriched in *cdc50*Δ EVs, whereas PI(34:2|16:0_18:2) (Fig 6E) were absent in *cdc50*Δ EVs. These observations suggest loss of Cdc50 reshapes the lipid cargo released by the *cdc50*Δ in addition to the increased EV production.

**Figure 6.**
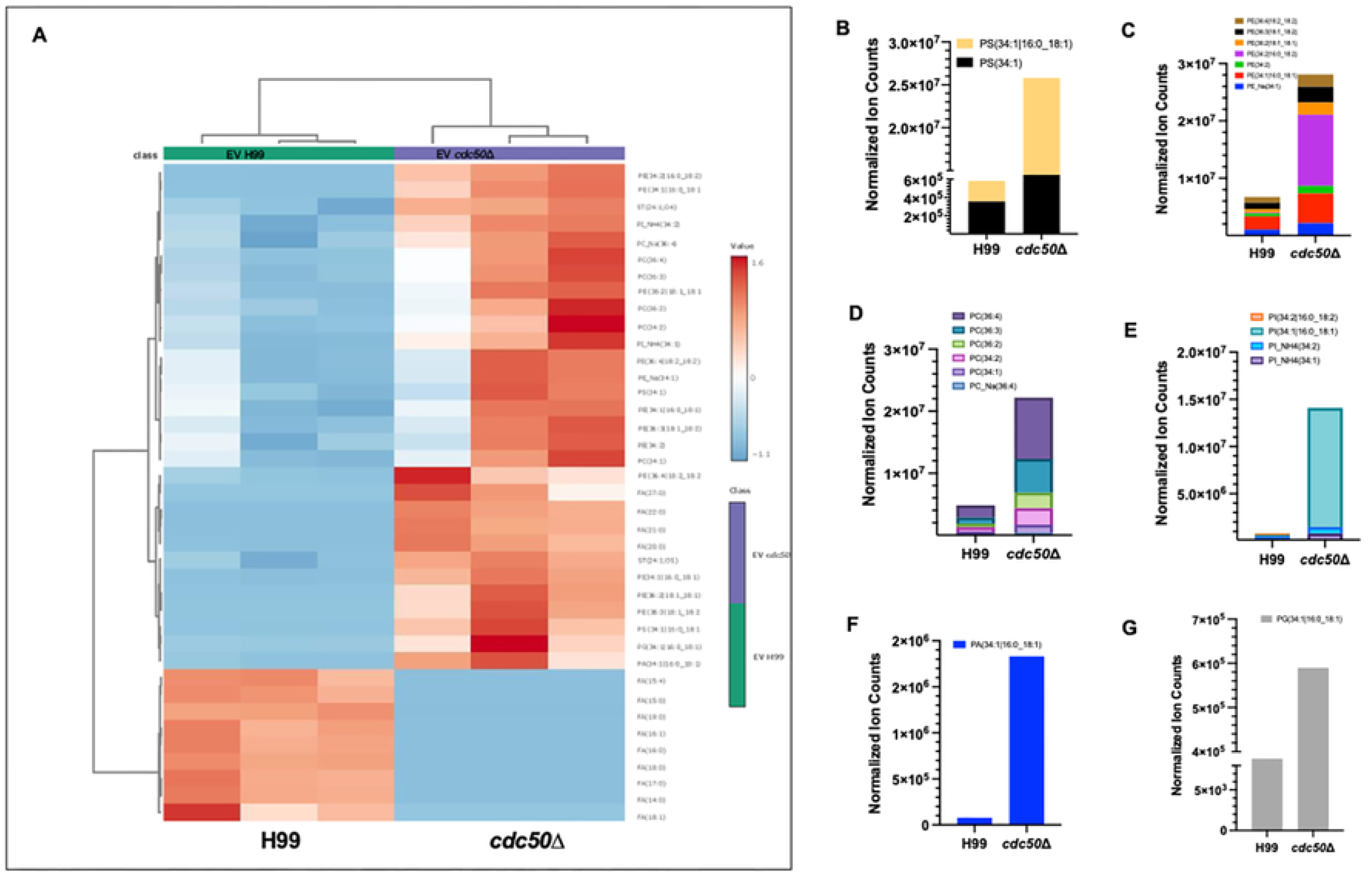
Lipid distribution in *cdc50*Δ and H99 EVs. (A) Heat map showing differentially expressed lipids in *C. neoformans* extracellular vesicles isolated from H99 and *cdc50*Δ (n=3 for each strain). The major classes of phospholipid species were presented: (B) Phosphatidylserine (PS); Phosphatidylcholine (PC); (C) Phosphatidylethanolamine (PE); (D) Phosphatidylcholine (PC); (E)Phosphatidylinositol (PI); (F) Phosphophatidic acid (PA) and (G) Phosphoglycerol (PG)

Furthermore, we also detected a reduction in fatty acid (FA) production in the *cdc50*Δ mutant EVs, primarily of FA below 20 carbon atoms compared to H99 EVs. Given that acyl-chain length influences membrane biology (29), we speculate these changes might have functional consequences for EV biogenesis and its interaction with host cells. Together, our results demonstrate loss of flippase function changes the lipid profile of both cryptococcal cells and their EVs, increasing PS exposure and enriching EVs for multiple phospholipid classes and species. This altered EV lipid composition may contribute to the distinct effects of H99 and *cdc50*Δ EVs on macrophage priming and fungal uptake.

### EVs differentially prime macrophages to modulate cryptococcal phagocytosis

Fungal EVs are known to interact with host cells and modulate immune responses (30–32). Given that the *cdc50*Δ mutant produced significantly more EVs with increased surface PS exposure while maintaining key vesicle architecture, we next asked whether these EVs could influence phagocytosis of different cryptococcal strains (9). Therefore, we hypothesized that EVs from H99 and *cdc50*Δ cells may differentially prime macrophages and alter subsequent fungal uptake. To test this hypothesis, we co-incubated J774 macrophage-like cells with EVs isolated from the *cdc50*Δ cells and infected with wild type H99 cells. Our results showed that the *cdc50*Δ EVs significantly increased the phagocytosis of H99 cells from 35% (no EV condition) to 50 % in average (Fig 7B). In contrast, when macrophages co-incubated with H99 EVs and infected with *cdc50*Δ cells significantly reduced the percentage of phagocytosis of *cdc50*Δ cells from 64% (no EV condition) to 41% (in the presence of H99 EVs) (Fig 7B), indicating strain dependent EV effect on phagocytosis.

**Figure 7.**
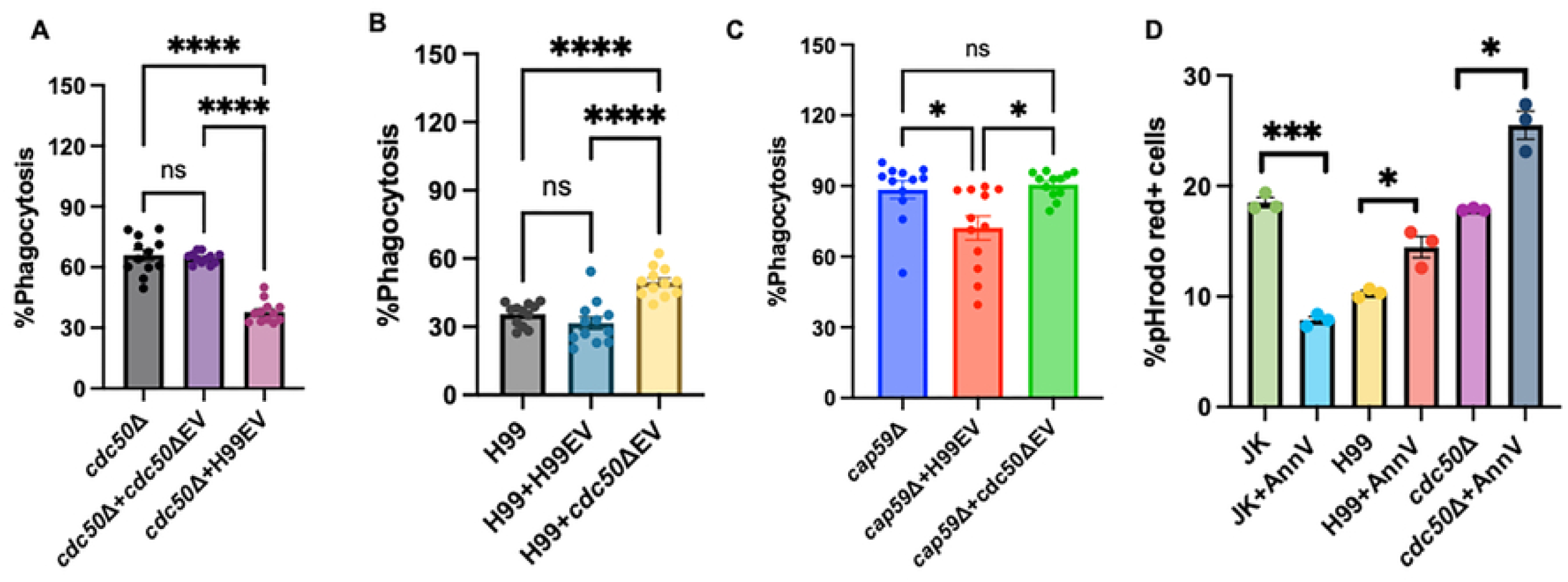
Extracellular Vesicles differentially prime macrophages to modulate cryptococcal phagocytosis. Macrophages were co-incubated with *cdc50*Δ EV or H99 EV and infected with either WT H99, *cdc50*Δ or *cap59*Δ for 2hr at 37℃ (A)*cdc50*ΔEV treated macrophages showed significant increase in % phagocytosis and yeast/macrophage ratio of WT cryptococcal cells compared H99EV treated macrophages infected with H99; (B) H99EV treated macrophages showed significant decrease in % phagocytosis and yeast/macrophage ratio of *cdc50*Δ cells compared to *cdc50*Δ EV infected with *cdc50*Δ; (C) Macrophages were either co-incubated with H99EV or *cdc50*ΔEV infected with *cap59*Δ, *cdc50*Δ EV treated macrophages showed significant increase % phagocytosis compared to H99EV stimulated macrophages. Bars represent mean± SEM. Statistical significance was determined using Welch’s one-way ANOVA followed by Dunnett’s T3 multiple-comparisons for panel A and B; and Kruskal-Wallis test followed by Dunn’s multiple-comparisons test for panel C. ns-not significant; ***p<0.0001 and *p<0.05

Given that H99 and *cdc50*Δ cells differ in capsule-associated phenotypes; because the cryptococcal capsule has anti-phagocytic function, we next tested whether EV-mediated modulation affects the phagocytosis of acapsular *cap59*Δ cells. We observed that J774 cells take up *cap59*Δ cells efficiently with over 90% phagocytosis rate, which is consistent with previous reports (33). A similar phenotype was observed for phagocytosis rate (over 90%) of J774 infected with *cap59*Δ and co-culturing in the presence of *cdc50Δ* EVs. Consistent with our findings, adding H99 EVs significantly decreased the phagocytosis of *cap59*Δ from 90% to 65 %, whereas *cdc50Δ* EVs did not significantly alter *cap59*Δ uptake (Fig 7C). Together, these results demonstrate that cryptococcal EVs are not only increased upon the loss of Cdc50 but can functionally prime macrophages to modulate cryptococcal phagocytosis.

### Cryptococcal uptake occurs independently of PS-MertK signaling

In apoptotic cells, the externalization of PS on the cells, sends out “eat me signal” to the macrophages, resulting in phagocytosis and allows the controlled elimination of these cells by the process of efferocytosis (15). Similarly, we observed that loss of Cdc50 results in elevated levels of PS exposure in fungal cell surface and increased phagocytosis on its interaction with macrophages (9). Therefore, we posit that macrophages recognize *cdc50*Δ better than the wild type H99 through PS-dependent efferocytosis pathways. Because EVs produced by *cdc50*Δ cells also had more surface PS, we further hypothesize that macrophages may recognize the PS signal of *cdc50*Δ cells through its secreted EVs.

To test this hypothesis, we blocked surface exposed PS with Annexin V prior to macrophage infection. Apoptotic Jurkat cells (JK cells) were included as a positive control for PS-dependent efferocytosis (34). Following the PS blockade, fungal and apoptotic cells were labelled with a pH sensitive dye pHrodo, and then co-cultured with J774 cells. Yeast cells engulfed by macrophages will turn red due to acidification of phagolysosome, which allows us to quantitatively assess phagocytic uptake using flow cytometry. Our data showed that Annexin V treatment significantly reduced macrophage uptake of apoptotic JK cells from 20% to 10%, confirming effective blockade of PS-mediated efferocytosis (Fig 7D). In contrast, PS blockade did not decrease the macrophage uptake of cryptococcal cells like that of JK cells. Instead, Annexin V treated cryptococcal cells showed increased uptake by J774 cells compared to untreated cryptococcal cells in both H99 (5% increase) and *cdc50*Δ (10% increase) strain backgrounds (Fig 7D).These results indicate although PS is exposed on the surface of *C. neoformans*, PS exposure might not be required for macrophage uptake of *C. neoformans.* The increased uptake observed after Annexin V treatment suggests that the exposed PS may not function as classic “eat-me” signal in this context and may instead participate in an inhibitory or masking interaction that limits fungal uptake. Alternatively, whether other phospholipids such as PE is required for host-*C. neoformans* interaction remain untested and warrants further investigation.

Because PS-mediated efferocytosis can occur through receptor-level signaling even when ligand availability is altered, we next examined whether macrophages nevertheless could recognize PS exposure in *cdc50*Δ cells through their canonical PS receptors. We first tested the potential involvement of PS receptor MertK, which belongs to Tyro3-Axl-MertK (TAM) receptor family (34). To test this directly, we employed mouse MertK γR1 (mMertKγR1) reporter cell line, in which activation of extracellular domain in the receptor protein triggers STAT1 phosphorylation in the presence of PS-binding molecule growth-arrest-specific 6 (Gas6) that can be detected by P-STAT1 antibody in a Western blot (Fig 8A) (34). The mMertKγR1 cells were incubated with multiple cryptococcal strains (H99, *cdc50*Δ, *cap59*Δ, and *cdc50*Δ *cap59*Δ), EVs from H99 and *cdc50*Δ, apoptotic JK cells, PS-expressing lymphoma cells (Cdc50L), and W3L cells (PS-negative control-lymphoma cells) for 30 mins before protein extraction, respectively (Fig 8B). In a western blot assay of extracted proteins from each treated mMertKγR1 cells, we observed robust STAT1 phosphorylation in the mMertKγR1 cells following exposure to apoptotic JK cells and Cdc50L cells, confirming the activation of Mertk signaling axis (Fig 8B) (34). In contrast, none of the cryptococcal strains or fungal EVs induced significant MertK activation signal intensity, even in the absence of capsule (Fig 8B). These results demonstrate that cryptococcal PS, whether presented on intact cells or on EVs, does not activate MertK receptor.

**Figure 8.**
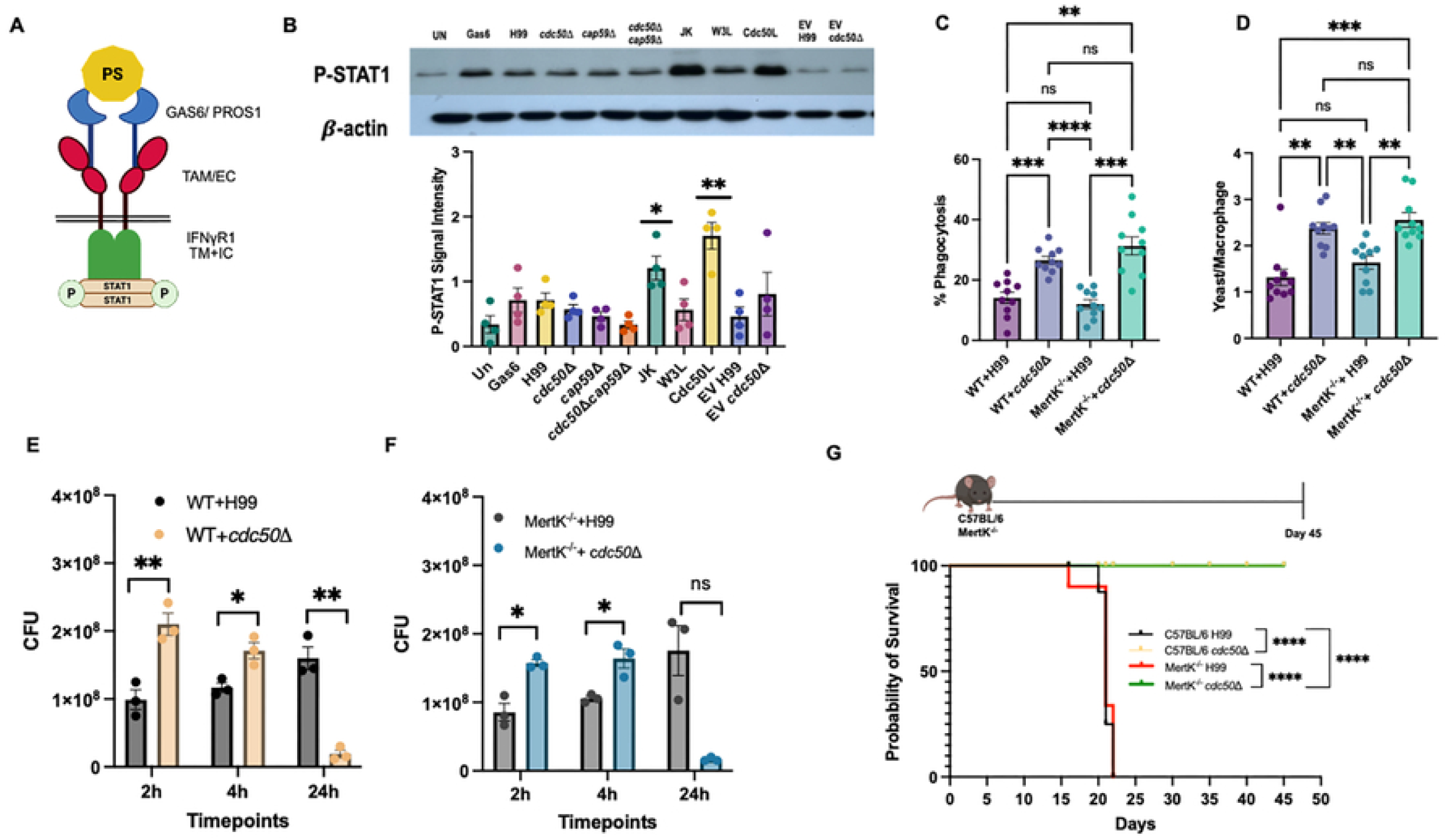
Cryptococcal phagocytosis occurs independent of PS receptor MertK. (A) Flow cytometry assay where J774 macrophages were infected with *C. neoformans* or apoptotic cells treated with 100 µg Annexin V and labelled with pHrodo. Percentage of pHrodo red+ cells define pHrodo labelled cells engulfed by macrophages defined by CD11b+F4/80+. Bars represent mean±SEM. Statistical significance was determined by a two-tailed unpaired t test with Welch’s correction (***p<0.0001 and * p<0.05). (B) Western blot analyses of phosphorylated STAT1 (pSTAT1) and b-actin in mMertKγR1 reporter cell line after treating reporter cell with different *C. neoformans* strains (H99, *cdc50*Δ, *cap59*Δ or *cap59*Δ *cdc50*Δ), cryptococcal EVs (*cdc50*Δ or H99), and apoptotic cells (JK) in presence of Gas6 for 30 mins. The level of MertKγR1 activation were measured by pSTAT1 signal intensities normalized to intensities of actin protein loading controls; (C-D) Activated macrophages were incubated with *C. neoformans* cells for 2 hr at 37℃ in 5% CO_2_. Following incubation, macrophages were washed, fixed with methanol, and stained with Giemsa. (C) The percentage of macrophages containing internalized yeast cells was determined by counting over 100 macrophages; (D) The yeast-to-macrophage ratio was quantified by inverted microscopy. Statistical analysis performed independently for each panel using Welch’s ANOVA followed by Dunnett’s T3 multiple-comparisons test, ns, not significant; **p<0.01; ***p<0.001; and p<0.0001. (E-F) Non-adherent extracellular yeast cells were then removed by washing with fresh DME. Number of fungal CFUs from macrophage cultures after an additional 0, 2, or 22 hr incubation were used to determine intracellular proliferation and macrophage killing in (E) WT and (F) MertK^-/-^. (G) Female C57BL/6 and MertK^-/-^ mice were infected with 1x 10^5^ yeast cells of either H99 or *cdc50*Δ strains. Survival curves for infected mice in a murine nasal inhalation model of systemic cryptococcosis. ****, Statistical significance by Mantel Cox test with Holm-Sidak multiple comparison p<0.0001.

To determine whether PS receptor MertK contributes to cryptococcal phagocytosis or fungal killing in vivo, we compared macrophage interaction with *C. neoformans* using WT and MertK^-/-^mice (35, 36). Our ex vivo macrophage interaction assay showed bone marrow derived macrophages (BMDM) isolated from both WT and MertK^-/-^ mice internalized *cdc50*Δ at significantly higher rates than H99 (Fig 8C-D). Consistent with this finding, *cdc50*Δ exhibited heightened sensitivity to macrophage-mediated killing in both WT and MertK^-/-^ mice (Fig 8E-F), indicating the increased phagocytosis during *cdc50*Δ-macrophage interaction is MertK receptor independent. Finally, to assess the potential contribution of MertK during infection, we infected both WT and MertK^-/-^ mice with either H99 or *cdc50*Δ. While H99 caused lethal infection in both mouse backgrounds at the similar rates, *cdc50*Δ was efficiently cleared in both mouse groups, with infected mice remaining healthy beyond 40 days post challenge (Fig 8G) and exhibited no fungal burden in the lungs and brains (**Supplemental Fig 1**). All these data indicate that loss of MertK did not alter host-*Cryptococcus* interaction, with either wild type or the *cdc50*Δ mutant. Thus, our data demonstrate that macrophage recognition, killing and in vivo clearance of *cdc50*Δ occurs independent of PS-MertK efferocytosis pathway. Instead, these findings support a model in which loss of Cdc50 results in a global membrane lipid dysregulation renders *C. neoformans* intrinsically susceptible to macrophage-killing, with PS exposure being a secondary consequence rather than a primary determinant of uptake.

### *cdc50*Δ significantly acidifies the macrophage phagosome

Having established that loss of Cdc50 causes broad lipid remodeling and ultrastructural defects in the plasma membrane, we next asked whether these membrane changes also alter the intracellular fate of *C. neoformans* inside the macrophages. Since the *cdc50*Δ phagocytosis occurs independently of PS-MertK signaling and Cdc50 has been reported to modulate Rim101 pathway (11), we hypothesized reduced survival of this mutant in the macrophages may reflect altered phagosomal acidification after its uptake by macrophages.

Since pHrodo fluoresces red in low pH environment, such as acidic compartments, to test our hypothesis, we labelled H99 and *cdc50*Δ cells with pHrodo dye prior to 2 h co-incubation with J774 cells, which allowed us to measure intracellular phagosomal acidification following fungal uptake by macrophages. Using flow cytometry, our results showed that macrophages containing *cdc50*Δ cells exhibited significantly higher pHrodo mean fluoresce intensity (over ∼75% increase) than macrophages containing H99 after phagocytosis (Fig 9A). This increase was also accompanied by a shift in fluorescence distribution (Fig 9B), indicating enhanced acidification of *cdc50*Δ-containing phagosomes compared with H99-conatining phagosomes, which leads to increased yeast killing.

**Figure 9.**
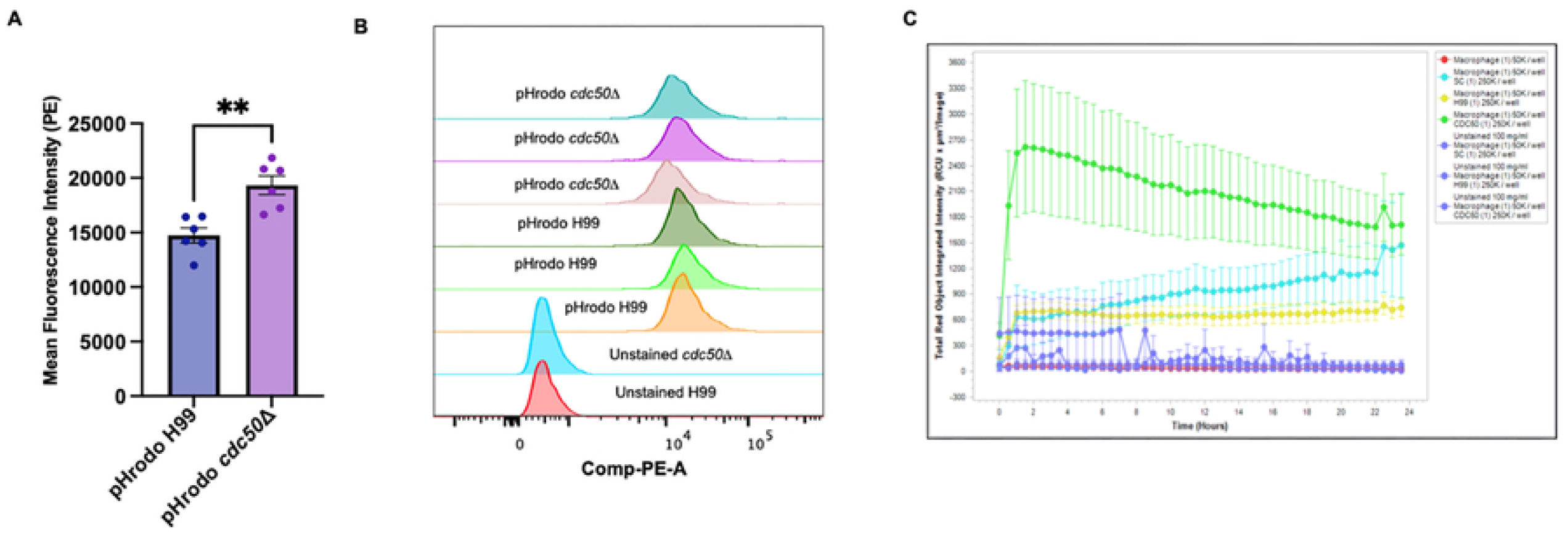
*cdc50*Δ significantly acidifies the macrophage phagosome. J774 macrophage like cells were co-incubated with pHrodo-labelled H99 or *cdc50*Δ cells for 2 hr (flow cytometry experiment) or 24 h for the Incucyte assay. (A) Fluorescent signal intensity was quantified using flow cytometry. (B) Representative images of fluorescent signal shifts using flow cytometry. (C) Cells were monitored in Incucyte Live-Cell Analysis System. Increasing numbers of labelled fungal strains in the presence of phagocytic cells yields increase in total red object area as fungal strains are engulfed. **Statistical significance was determined using Unpaired two tailed t test with Welch’s correction p<0.05

To further examine the kinetics of this response, we monitored pHrodo fluorescence over 24h using an Incucyte live-cell analysis system (Sartorius, New Jersey). This platform enables a dynamic measurement of acidification kinetics rather than a single endpoint readout, allowing us to track real time changes in pHrodo fluorescence following macrophage-fungal interactions and compare the progression of acidification between strains. Consistent with our flow data, we showed that macrophages co-incubated with *cdc50*Δ cells triggered a rapid and substantial increase in acidification signal, whereas H99-containing compartments remained less acidified across the time course (Fig 9C). Together, these results demonstrate that loss of Cdc50 enhances phagosomal acidification following macrophage uptake. These findings suggest that the membrane defects caused by loss of Cdc50 not only affect fungal recognition and uptake but also compromises its ability to withstand intracellular phagosomal processing, providing a potential explanation for increased susceptibility of *cdc50*Δ cells to macrophage-mediated killing.

## Discussion

Successful establishment of cryptococcosis depends on the ability of *C. neoformans* to coordinate cell surface structure organization, vesicular trafficking, and immune evasion during its encounter with host macrophages. While fungal extracellular vesicles (EVs) are increasingly recognized as important mediators of host-pathogen communication, the lipid regulators that shape EV biogenesis, lipid composition, immune response remain poorly defined. In this study, we identified cryptococcal Cdc50 as a key player in membrane lipid homeostasis that regulates EV output and modulates macrophage responses. By integrating lipidomics, ultrastructural analysis, EV biology, and host-pathogen interaction assays, our findings revealed how the disruption of phospholipid asymmetry in *cdc50*Δ rewires fungal plasma membrane lipid dynamics, leading to altered macrophage recognition and macrophage-mediated clearance of fungal cells.

Cdc50 is the regulatory subunit of phospholipid translocase (flippase), consisting of two transmembrane domains and a large luminal loop. By interacting with the catalytic subunit P4-ATPase, Cdc50 is essential for regulating phospholipids translocation to maintain plasma membrane lipid asymmetry, a fundamental determinant of membrane organization and cellular homeostasis (6, 27). Our previous study showed loss of Cdc50 function in *C. neoformans* abolished the flippase activity, disrupted lipid asymmetry resulting in increased PS accumulation on its cell membrane surface (9). We speculate that disruption of phospholipid asymmetry in the flippase mutant lowers energetic barrier for membrane deformation, thereby favoring increased outward vesiculation from the plasma membrane. Importantly, hyperproduction of EVs in *cdc50*Δ cells may have occurred also due to severe membrane destabilization, which would suggest that loss of lipid asymmetry drives excessive vesiculation through compromised membrane architecture rather than regulated budding or nonspecific cell lysis.

Accumulating evidence indicates that fungal EVs are biologically active structures that engage host immune cells and modulate macrophage behavior during infection (19, 20). Fungal EVs are packed with nucleic acids, proteins, lipids, polysaccharides, glycans, small molecules and small metabolites (37, 38), aid in intraspecies and cross-species communication (24, 39). In *Cryptococcus*, EVs have previously been shown to influence macrophage uptake. For instance, EVs derived from acapsular *cap67Δ* mutant cells were able to enhance the phagocytosis of encapsulated yeast (33). Because *cdc50*Δ produced increased numbers of EVs, we tested whether these vesicles are functionally active during macrophage interaction. In our study, EVs were presented simultaneously with fungal cells during macrophage interaction, allowing EVs to influence host-pathogen interactions in real time rather than through prior macrophage conditioning. Under these conditions, EVs exerted strain dependent effects on phagocytosis, in which *cdc50*Δ EVs promoted phagocytosis whereas H99 EVs reduced phagocytosis. Notably, coincubation of EVs with acapsular *cap59*Δ yeast recapitulated this trend. These results demonstrate that the differential lipid composition of the EVs actively shape early macrophage-*C. neoformans* interaction thereby altering the macrophage response. Overall, the cryptococcal EVs function as dynamic immunomodulatory cues that influence cryptococcal uptake by macrophages in a strain specific manner, *cdc50*Δ EVs enhance phagocytosis whereas H99 EVs promotes suppression, possibly due to difference in their surface pathogen associated molecular patterns (PAMPs), which warrants further studies.

In fungi, multiple components governing plasma membrane architecture and lipid metabolism have been directly implicated in EV biogenesis and cargo specification. For instance, in *Candida albicans* disruption of phospholipid biosynthesis impacted EV morphology and composition of the cargo (40). In *C. neoformans,* deletion of Apt1 resulted in EVs with a lower GXM and polysaccharide concentration (41, 42). Thus, the distinct effects of H99 and *cdc50*Δ EVs on macrophage uptake may be linked to differences in the EV lipid composition. *cdc50*Δ EVs are enriched in phospholipids and exhibited reduced abundance of fatty acid species, particularly those with fewer than 18 carbon atoms, suggesting that EVs preferentially incorporate long-chain lipids to enhance bilayer thickness and stability in extracellular environment (43). Together, these findings indicate that EVs are not simple reflections of cellular lipid composition but represent selectively sorted lipid entities released by the cells. Altogether, these findings strongly support the hypothesis that plasma membrane is a fundamental and dynamic cellular site for fungal EV biogenesis, where plasma membrane lipid asymmetry might be contributing to its cargo selection. Moreover, how do vesicles select specific components for export is yet to be deciphered.

The outer membrane leaflet consists of phosphatidylcholine (PC) and sphingomyelin whereas the inner membrane consists of PS and PE. In the mammalian cell, redistribution of PS to the outer membrane flags the cell for apoptosis and is cleared by macrophage-mediated efferocytosis through the engagement of multiple PS-sensing receptors including the TAM (Tyro3, Axl and MertK) receptor family (15) and CD300 family (16–18). In our previous studies, we found that the expression of *CDC50* was upregulated during *C. neoformans*-macrophage interaction (9) and the *cdc50*Δ mutant showed PS accumulation on plasma membrane surface and increased susceptibility to macrophage mediated killing (9), an observation similar to the PS triggered efferocytosis. This similarity leads to our hypothesis that elevated PS exposure on *cdc50Δ* cells and EVs might also serve as a recognition signal for macrophages to take up the mutant cells (9). However, our current study revealed that cryptococcal PS did not engage in canonical PS recognition pathways. In our study, PS masking using Annexin V did not affect the cryptococcal uptake by macrophages, interestingly, we observed an increased phagocytosis after Annexin V addition. We speculate Annexin V binding may modify the fungal surface or relieve a PS-associated inhibitory interaction, allowing macrophages to recognize the other cell wall or membrane-associated ligands more efficiently. It is also possible that blocking PS may induce the surface presentation of other lipids that could alter the host-*Cryptococcus* interaction. Therefore, PS exposure in *cdc50Δ* can be interpreted as a sign of disrupted membrane asymmetry rather than primary ligand driving phagocytosis. Because PS-mediated efferocytosis can occur through receptor-level signaling even when ligand availability is altered (15), we next assessed activation of high affinity PS receptor MertK using highly sensitive MertKγR1 reporter system (34, 44, 45). While apoptotic Jurkat cells (JK) and PS-expressing Cdc50L cells robustly activated MertK, neither cryptococcal cells nor cryptococcal EV elicited MertK activation, irrespective of capsule status. Consistent with these findings, macrophage uptake, intracellular killing, and in vivo clearance of *cdc50*Δ cells were observed in MertK^-/-^ mice at a similar level as that of the naïve mice. Moreover, we performed qRT-PCR but did not detect an upregulation of other PS receptor expression, including Axl, Tyro3 and TIM family receptors, in either J774 cell line or primary BMDM cells when co-incubated with *cdc50*Δ cells (**Supplemental figure 2-3**). Although CD300f receptor has been reported to sense PS in pathogenic and non-pathogenic *Rickettsia* species (46), in our qRT-PCR assay (**Supplemental figure 4, Table S1**), PS accumulation on the *cdc50*Δ did not induce the expression of CD300 receptors. All these data suggest that PS-PS receptor recognition may not be the mechanism of increased phagocytosis in *cdc50*Δ-macrophage interaction. However, it remains possible that other PS receptors, and other fungal phospholipid species may be involved in *Cryptococcus*-host interaction. In the future, we could test other PS receptor report cell lines, additional receptor KO mice, or using receptor inhibitors, but those works will be beyond the scope of this study.Together, these results demonstrate that PS exposure on *C. neoformans*, whether presented on whole cells or EVs, is not functionally equivalent to mammalian PS, hence, not recognized via MertK receptor or other PS receptors we analyzed. Moreover, our results demonstrate PS externalization in *cdc50Δ* represents a downstream consequence of global membrane lipid dysregulation rather than a primary immunological signal.

The lack of PS-MertK dependence led us to consider whether increased uptake and killing reflect a broader membrane-homeostasis defect rather than a single PS receptor-ligand interaction. Our previous study showed loss of Cdc50 function in *C. neoformans* abolished the flippase activity, leading to increased PS accumulation on its cell membrane surface. However, how loss of Cdc50 affects fungal membrane dynamics remains unknown. Using whole cell lipidomic analysis, we detected an increased accumulation of phospholipids, including PS, and reduced fatty acid (FA) levels, indicating both an altered phospholipid distribution and a global remodeling of membrane lipids in *cdc50*Δ. This result is consistent with a previous report by Brown et al. (11) Importantly, increase in PA and PS promote negative membrane curvature (47, 48), suggesting their enrichment may facilitate vesicle budding in *cdc50*Δ. While our results showed a broad reduction in multiple saturated fatty acid production, we also observed increase in FA(18:1), sterol ST(24:1:O5) and ST(24:1:O4). Fatty acid synthesis is essential for *C. neoformans* survival (49). Together, our findings suggest that disruption of lipid asymmetry may perturb fatty acid synthesis, where increase in monounsaturated fatty acid and sterol might serve as a compensatory mechanism to restore lipid order and integrity, which in turn also increases the susceptibility of *cdc50*Δ to antifungal drugs (9, 10, 49). Concomitant with these compositional perturbances, *cdc50*Δ cells exhibited pronounced ultrastructural defects like plasma membrane invaginations resulting in aberrant intracellular radiopaque structures in the cytoplasm, indicating that the increased phospholipids in the mutant might have resulted in an excess or mismanaged membrane synthesis.

Alveolar macrophages are the first line of defense against *Cryptococcus* infection. Following the phagocytic uptake, *C. neoformans* is contained inside the phagosome that matures to form phagolysosome, a hostile compartment characterized by progressive acidification for the destruction of the pathogen inside (50, 51). However, the glucuronic acid present in capsular polysaccharide helps to buffer the phagosomal acidification (52). Fungal cells can replicate inside the phagosome and dictate the composition of phagosomal compartment in a process associated with the formation of a leaky phagosome and accumulation of cytoplasmic vesicles filled with capsular polysaccharide, which also aid their escape via non-lytic vomocytosis (53–57). Overall, the interaction of macrophages with *C. neoformans* results in either dissemination, latency or clearance of yeasts (4, 54). Our previous study showed loss of Cdc50 resulted in capsule enlargement yet the mutant cells being rapidly cleared by the macrophages. Having established that loss of Cdc50 causes broad membrane lipid remodeling and structural defects, we next considered whether these changes alter the intracellular fate inside the macrophage. By monitoring phagosomal acidification in real time using pH sensitive dye pHrodo, we detected a more rapid and sustained increase in acidification signal over time in the *cdc50*Δ cells than the macrophages containing WT H99. These findings are in line with the previous studies demonstrating that *C. neoformans* alters phagosomal maturation to create permissive intracellular niche for its survival (53, 58–60). In contrast, loss of Cdc50 compromises the ability of *C. neoformans* to resist phagosomal acidification, providing a mechanistic explanation for heightened susceptibility of *cdc50*Δ cells to macrophage mediated killing.

Beyond its role in membrane biology, the flippase complex has emerged as a promising antifungal target. Although *C. neoformans* is intrinsically resistant to echinocandin drug caspofungin, disruption of flippase function increased its susceptibility to echinocandins through hyperactivation of calcineurin pathway (9, 61, 62). Further strengthening the therapeutic potential of targeting lipid flippase, we designed antifungal peptides targeting flippase function in *C. neoformans* by blocking the Cdc50-P4-ATPase interaction and identified a potent peptide-based inhibitor Cryptomycin KKOO (63, 64). The peptide treatment sensitized *C. neoformans* wild type H99 to phenotypes of the *cdc50*Δ mutant, including increased sensitivity to anti-fungal drugs, macrophage killing, increased reactive oxygen species and intracellular calcium levels. These data indicated that the peptide targeted flippase function (63, 64). Furthermore, a recent study identified a nature product butyrolactol A as a fungal lipid flippase inhibitor that sensitized echinocandin drugs (65). Mechanistic study revealed that butyrolactol A trapped the flippase Cdc50-Apt1 complex in a non-functional state, leading to disrupted membrane asymmetry, impaired vesicular trafficking and cytoskeletal organization, thereby enhancing the uptake and antifungal efficacy of echinocandins (65). Together, these findings highlight membrane lipid asymmetry as an attractive antifungal target.

Based on our findings, we propose a model in which Cdc50-dependent lipid asymmetry functions as a regulatory checkpoint linking membrane homeostasis to macrophage recognition and phagocytosis in *C. neoformans*. In WT cells, membrane lipid homeostasis maintains membrane asymmetry and architecture, and produces EVs that suppress macrophage phagocytosis, aiding in fungal survival and dissemination. In contrast, loss of Cdc50 leads to global membrane lipid dysregulation resulting ultrastructural membrane defects, and increased production of phospholipids enriched EVs that enhanced macrophage uptake. This enhanced macrophage phagocytic activity is independent of PS mediated efferocytosis. Following the uptake, *cdc50*Δ cells fail to resist phagosomal acidification, rendering them intrinsically susceptible to macrophage mediated killing and *in vivo* clearance Thus, Cdc50 emerges as a key regulator of membrane lipid homeostasis that influences host-pathogen interactions.

## MATERIALS AND METHODS

### Strains and Media

*Cryptococcus neoformans* clinical strain H99 and its mutants *cdc50*Δ, complement strain (*cdc50Δ+ CDC50*), as well the *cap59*Δ, and *cdc50Δ cap59Δ* double mutant used in this study were cultured in Yeast extract-peptone-dextrose (YPD) medium at 30°C with moderate shaking overnight (or otherwise specified) for the experiments.

### Cell Culture

Macrophage-like cell line J774 cells were cultured in DME medium with 10% heat-inactivated FBS at 37°C with 5% CO_2_. mMertKYR1 cells were cultured in HAM’s Media supplemented with 10% heat-inactivated FBS, 5 % Penstrep and L-glutamine at 37°C with 5% CO_2_. Lymphoma cell lines W3L and Cdc50L were cultured in RPMI media supplemented with 10% heat-inactivated FBS, 5% Penstrep and L-glutamine at 37°C with 5% CO_2_

### Lipidomic analysis

Briefly, for lipid isolation from *C. neoformans*, the cells were grown overnight in YPD liquid media. Next day, the cells were washed three times and set to the concentration of 1×10^8^ cells/ml. For lipid isolation from *C. neoformans* EVs, the EVs were dried by placing the tube upside down for several hours. For lipid extraction from the cells and EVs, 5 volumes of −20°C chloroform-methanol (2:1) were added with SPLASH and incubated on ice for 5 mins, centrifuged for 10 mins at 4°C at 12,000 rpm. The bottom lipid phase was collected in amber colored autosampler tubes. The samples were dried out using steady stream of nitrogen gas. Lipids were dissolved in methanol and analyzed by liquid chromatography-tandem mass spectrometry (LC-MS/MS) in a Q-Exacative mass spectrometer (ThermoFischer, Waltham, MA). Lipid species were identified with SPLASH and manually inspected before having the features (peak areas) extracted with MZmine 2.0. Samples were normalized to internal standard and the heatmaps were plotted using Metaboanalyst software (https://www.metaboanalyst.ca/MetaboAnalyst/).

### Transmission electron microscopy

For morphological analyses at the ultrastructural level, cells were fixed in 2% paraformaldehyde/2.5% glutaraldehyde (Ted Pella Inc., Redding, CA) in 100 mM cacodylate buffer, pH 7.2 for 2 hr at room temperature and then overnight at 4°C. Samples were washed in cacodylate buffer and postfixed in 1% osmium tetroxide (Ted Pella Inc.)/ 1.5% potassium ferricyanide (Sigma, St. Louis, MO) for 1 hr. Samples were then rinsed extensively in dH_2_O prior to en bloc staining with 1% aqueous uranyl acetate (Ted Pella Inc.) for 1 hr. Following several rinses in dH_2_O, samples were dehydrated in a graded series of ethanol and embedded in Eponate 12 resin (Ted Pella Inc.). Ultrathin sections of 95 nm were cut with a Leica Ultracut UCT ultramicrotome (Leica Microsystems Inc., Bannockburn, IL), stained with uranyl acetate and lead citrate, and viewed on a JEOL 1200 EX transmission electron microscope (JEOL USA Inc., Peabody, MA) equipped with an AMT 8 megapixel digital camera and AMT Image Capture Engine V602 software (Advanced Microscopy Techniques, Woburn, MA).

### Extracellular Vesicles Isolation

Cryptococcal EVs were isolated according to a previously described solid media method by Reis et al 2019, with some modifications. In brief, loop full of cells were inoculated in 5 mL YPD media 30°C overnight with shaking 225 rpm. Next day, the cells were washed twice with 10 mL of sterile water, set to a density of 3.5 × 10^7^ cells/ml in water. Aliquots of 300 μl of cell suspensions were spread onto YPD plates and incubated for 24 hr at 30°C. The cells were carefully recovered into 10 mL of 1x 0.22 μm-filter sterile PBS. Pellet by centrifugation at 5000 x g for 15 mins at 4°C. The supernatant was again centrifuged at 10,000 x g for 30 mins at 4°C. The supernatant was filtered using 0.45 μm syringe filter and ultracentrifuged ay 100,000 x g for 2 hr at 4°C (Ti70 bucket rotor, Beckman coulter) in ultra clear tubes. The supernatant was discarded and the pellet was resuspended in 0.22 μm pore-filtered PBS. The pellets were stored at −80°C until use or subjected to NTA (by diluting in sterile water or PBS) for measuring concentration and diameter. (66)

### Cryo-EM for EVs

EVs (4 μl) were spotted on Quantifoil holey carbon grids, R1.2/1.3 and cryo-fixed by plunge freezing in liquid nitrogen using ThermoFisher Vitrobot mark IV. Grids were observed with ThermoFisher Tundra 100 TEM operating at 100 keV using the software EPU (ThermoFisher Scientific) for data collection.

### Nanoparticle tracking analysis (NTA) of EVs

This method was used for monitoring the presence of EVs in our samples (NTA). This analysis was performed on an LM10 nanoparticle analysis system, coupled with a 488-nm laser and equipped with an SCMOS camera and a syringe pump (Malvern Panalytical, Malvern, UK). All samples were measured as described above. The data were acquired and analyzed using the NTA 3.0 software (Malvern Panalytical). Particle numbers were obtained by NTA. NTA histograms were also used for the quantification of EV peaks in different samples. EV diameters were also measured using NTA. (67)

### Flow Sorting of Extracellular Vesicles

The vesicles were diluted to a concentration of 1 x 10^8^ particles/ml in 1x Annexin V binding buffer (10x Annexin V binding buffer recipe: 0.1M Hepes (pH 7.4), 1.4 M NaCl, 25mM CaCl_2_). Vesicles were stained with 2-dialkylcarbocyanine dye namely DiL dye (1,1-dioctadecyl-3,3,3,3-tetramethylindocarbocyanine perchlorate; Vybrant DiI cell labeling solution) following (68) with some modifications. Briefly, the EVs were stained with DiL dye for 1 hr at 4°C followed by ultracentrifugation for 2 hr at 100,000 x g 4°C (Ti70 bucket rotor Beckman Coulter). The vesicles were then resuspended in 1x Annexin V binding buffer and incubated at room temperature for 15-20 mins in dark with 5 μl of Annexin V binding buffer. Dye-stained vesicles were analyzed with a BD FACS cell sorter. Data was analyzed using FLowJo software.

### BMDM isolation

Bone marrow-derived macrophages (BMDM) were isolated from the tibia and femurs of 8–12-week-old C57BL/6 WT mice or MerTK^-/-^ mice. The tibia and femurs were isolated and washed for 15 seconds each, once in 70% ethanol and then in PBS, following which their ends were cut to reveal the bone marrow. The bones were placed cut end facing down in a 0.5 ml Eppendorf tube with a hole pierced with an 18 G needle, which was placed in a 1.5 ml Eppendorf tube containing 80 μl of BMDM media. The tubes were then centrifuged at 15,000×*g* to allow the bone marrow to precipitate into the 1.5 ml Eppendorf tube. The cell pellet was treated with 5 ml of RBC lysis buffer for 5 min at room temperature, later quenching with BMDM media. The cells were collected by centrifuging at 300×*g* for 5 min and strained using a 70-micron cell strainer. The cells were counted using a hemocytometer and were plated in cell culture-treated plates at 10^7^ viable cells per plate, in a total volume of 30 ml. Recombinant murine M-CSF (Biolegend-576406) was added at a final concentration of 20 ng/ml. On day 3 of differentiation, 15 ml of the cell culture medium was removed and replaced with 15 ml of fresh M-CSF-containing medium. BMDMs were fully differentiated on day 7. Differentiated BMDMs were detached gently using a scraper and re-plated into 12-well, 24-well or 48-well plates based on experimental needs.

### *Cryptococcus*-macrophage interaction assay

J774 cells (5 × 10^4^) or BMDM in 0.5 ml fresh DME medium were added into each well of a 48-well culture plate and incubated at 37°C in 5% CO_2_ overnight. To activate macrophage cells, 50 units/ml gamma interferon (IFN-γ; Invitrogen) and 1 µg/ml lipopolysaccharide (LPS; Sigma) were added into each well. *C. neoformans* overnight cultures were washed with phosphate-buffered saline (PBS) twice and opsonized with 20% mouse complement. *Cryptococcus* cells (2 × 10^5^) were added into each well (yeast/J774 ratio, 4:1) To assess the effect of EVs on phagocytosis the *Cryptococcus* cells were either cocultured with H99 EVs or cdc50Δ EVs at concentration of 10^8^ particles/ml. To assess the phagocytosis rate, the cells were washed with PBS after a 2-hr coincubation and fixed with methanol for 30 min. Giemsa stain was added to the wells at a 1:10 dilution, and the plates were incubated overnight at 4°C. Cells were washed once with PBS and analyzed using an inverted microscope. For each well, 3-5 different fields were counted, for a total of at least 100 macrophages. The percent phagocytosis was determined by dividing the number of macrophages that contained *C. neoformans* by the total number of macrophages counted. (9)

### Macrophage Killing Assay

BMDM cells (5 × 10^4^) in 0.5 ml fresh DME medium were added into each well of a 48-well culture plate and incubated at 37°C in 5% CO_2_ overnight. To activate macrophage cells, 50 units/ml gamma interferon (IFN-γ; Invitrogen) and 1 µg/ml lipopolysaccharide (LPS; Sigma) were added into each well. *C. neoformans* overnight cultures were washed with phosphate-buffered saline (PBS) twice and opsonized with 20% mouse complement. *Cryptococcus* cells (2 × 10^5^) were added into each well (yeast:J774 ratio, 4:1). To assess intracellular proliferation of *C. neoformans*, nonadherent extracellular yeast cells were removed by washing with fresh DME medium after a 2-hr coincubation and cultures were incubated for another 0, 2, and 22 hr. At indicated time points, the medium in each well was replaced with distilled water (dH_2_O) to lyse macrophage cells for 30 min at room temperature. The lysate was spread on YPD plates, and CFU were counted to determine intracellular proliferation. (9)

### mMertKγR1 Binding Assay

Briefly, this reporter cell line contains mouse MertK extracellular domain and transmembrane domain and intracellular domains of STAT-1. Ligand induced binding results in pSTAT1 activation. For the binding assay, 1×10^6^ cells were plated onto 6-well plate, serum starved for at least 6 hr. in serum-free HAM’s F12 media. Cells were incubated with HEK_293_TN cell conditioned media containing GAS6 for 5 mins and then cells were treated with apoptotic cells, PS lipid, cryptococcal cells or cryptococcal EVs for 30 mins at 37°C in 5% CO_2_. Next, cells were washed with ice cold PBS thrice and were lysed with HNTG lysis buffer (20mM HEPES, pH 7.5, 150 mM NaCl, 10% Glycerol) supplemented with 10% triton X-100, 1 mM PMSF, 1 mM sodium orthovanadate, 10 mM sodium molybdate, 1 mM EDTA and 1% protease inhibitor cocktail (Thermo Fisher)). Whole cell lysates were subjected to Western blot to determine the activation of mMertKγR1 cell line.

### Western Blot

Whole cell lysates were prepared in HNTG lysis buffer. Bradford assay was done to measure protein concentration and 5 µg of protein for each sample was loaded onto a SDS reducing gel (10% acrylamide). SDS gel electrophoresis was carried out at 60 V and then at 120 V when proteins entered the resolving gel. Proteins were transferred from the gel onto a PVDF membrane using the wet transfer method at a constant current of 0.25 A for 70 mins. Blots were blocked in 5% BSA made in TBST overnight. Blots were incubated 1 hr or overnight with primary antibodies, anti-PSTAT-1 (Cell signaling Technologies) and anti-β-ACTIN (Cell signaling Technologies) at 1:1000 dilution at 4°C. Next, blots were washed thrice with TBST for intervals of 15 mins. The blots were then incubated with secondary antibodies, anti-mouse IgG-HRP (1:4000 anti-PSTAST-1 and 1:10,000 for anti-β-ACTIN) for 1 hr. at room temperature. After 1 hr, the blots were washed thrice with TBST for intervals of 15 minutes. The blots were developed with ECL substrate (Bio-Rad (Hercules, CA)) and chemiluminescent images were captured using Bio-Rad Image Doc (Hercules, CA). The level of MertKγR1 activation were measured by pSTAT1 signal intensities normalized to intensities of actin protein loading controls.

### Efferocytosis Assay

J774 Macrophage like cells were plated onto 24-well plate and cultured in DME medium containing 10% HI-FBS. Apoptotic cells were generated by treating Jurkat cells with 1 µM Staurosporine (FUJIFILM) for 3hr at 37 °C in RPMI medium without serum. All the target cells (apoptotic cells & cryptococcal cells) were labelled with pHrodo 125 ng/ml for 1 hr in dark at 37 °C. Next, the cells were washed three times with PBS. The labelled target cells were added to plated macrophages at a at 5:1 ratio activated using above cryptococcus macrophage interaction protocol. The co-culture was incubated for 2hr at 37 °C. After incubation, macrophages were washed twice with PBS and detached by trypsinization. Efferocytosis was assessed by flow cytometry, measuring the percentage of pHrodo^+^ cells within the CD11b^+^ F4/80^+^ macrophage population.

### Incucyte Assay

J774 Macrophage like cells were plated onto 24-well plate and cultured in DME medium containing 10% HI-FBS. Apoptotic cells were generated by treating Jurkat cells with 1 µM Staurosporine (FUJIFILM) for 3 hr at 37 °C in RPMI medium without serum. All the target cells (apoptotic cells & cryptococcal cells) were labelled with pHrodo 125 ng/ml for 1 hr. in dark at 37 °C. Next, the cells were washed three times with PBS. The labelled target cells were added to plated macrophages at a 5:1 ratio activated using above cryptococcus macrophage interaction protocol. The co-culture was incubated for 24 hr at 37 °C in Sartorius Incucyte machine for real time phagosomal acidification kinetics. Data was analyzed using Sartorius analysis software.

### Murine infection assays

Groups of 8 female C57/B6 mice and MertK^-/-^ were intranasally infected with 10^5^ yeast cells of each strain as previously described in (9). Over the course of the experiments, animals that appeared moribund or in pain were sacrificed by CO_2_ inhalation. Survival data from the murine experiments were statistically analyzed using the log rank test of the Prism (GraphPad Software, San Diego, CA). *P* values of <0.001 were considered significant. Infected animals were sacrificed at the endpoint of the experiment according to the Rutgers IACUC-approved animal protocol. For mice infected by the *cdc50*Δ mutant strain, the experiment was terminated 45 days postinfection. To compare the survival, lungs, spleens and brains from mice infected by H99 and the *cdc50*Δ mutant, were isolated at endpoint. Infected organs were homogenized in 1xPBS buffer using a homogenizer. Resuspensions were diluted, and 100 µl of each dilution was spread on YPD medium; fungal colonies were counted after 3 days of incubation at 30°C.

## Statistical Analysis

GraphPad Prism software was used to analyze the data statistically. All experiments were repeated at least three times. Experiments were examined for statistical significance using a two-tailed Student *t* test and One-way ANOVA to compare data from multiple groups*. P* <0.05 were considered significant.

## Ethics Statement for Animal Use

Animal studies were performed at Rutgers University Newark campus animal facility. All studies were conducted following biosafety level 2 (BSL-2) protocols and procedures approved by the Institutional Animal Care and Use Committee (IACUC) and Institutional Biosafety Committee of Rutgers University under protocol 999901066. Animal studies were compliant with all applicable provisions established by the Animal Welfare Act and the Public Health Services (PHS) Policy on the Humane Care and Use of Laboratory Animals.

## Biorender License

Schematic figures are made using Biorender (https://app.biorender.com).

## Author contributions

Siddhi Pawar Data curation, Formal analysis, Investigation, Methodology, Writing-original draft, Writing | Yu Zhang, Data Curation, Methodology | Christopher Varsanyi Formal analysis, Methodology | Varsha Gadiyar Formal analysis, Methodology | Samantha Avina, Formal analysis, Methodology | Raymond Birge Methodology, Resources and Supervision | Chaoyang Xue, Conceptualization, Formal analysis, Supervision, Funding acquisition, Project administration, resources, writing – review and editing.

## Acknowledgements

We thank all members of Xue lab and Birge lab for their support. We also thank Tamara Doering’s lab and Wendy Beatty at Washington University School of Medicine, St louis for technique support and TEM Imaging. We also thank Amariliz Rivera and Kabilan Velliyagounder for helpful discussions to better these studies. This project is supported by NIH grant AI155647 and AI141368 to CX, and NCI 5R01CA260137 to RB.

## Reference

1. Rajasingham R, Smith RM, Park BJ, Jarvis JN, Govender NP, Chiller TM, Denning DW, Loyse A, Boulware DR. 2017. Global burden of disease of HIV-associated cryptococcal meningitis: an updated analysis. Lancet Infect Dis 17:873–881.

2. Coelho C, Bocca AL, Casadevall A. 2014. The intracellular life of *Cryptococcus neoformans*. Annu Rev Pathol 9:219–38.

3. Lin X, Heitman J. 2006. The biology of the *Cryptococcus neoformans* species complex. Annu Rev Microbiol 60:69–105.

4. Wang Y, Pawar S, Dutta O, Wang K, Rivera A, Xue C. 2022. Macrophage Mediated Immunomodulation During *Cryptococcus* Pulmonary Infection. Frontiers in Cellular and Infection Microbiology Volume 12–2022.

5. Pirofski L-a, Casadevall A. 2017. Immune-Mediated Damage Completes the Parabola: *Cryptococcus neoformans* Pathogenesis Can Reflect the Outcome of a Weak or Strong Immune Response. mBio 8:10.1128/mbio.02063-17.

6. Tanaka K, Fujimura-Kamada K, Yamamoto T. 2011. Functions of phospholipid flippases. J Biochem 149:131–43.

7. Stanchev LD, Rizzo J, Peschel R, Pazurek LA, Bredegaard L, Veit S, Laerbusch S, Rodrigues ML, López-Marqués RL, Günther Pomorski T. 2021. P-Type ATPase Apt1 of the Fungal Pathogen Cryptococcus neoformans Is a Lipid Flippase of Broad Substrate Specificity. J Fungi (Basel) 7.

8. Veit S, Laerbusch S, López-Marqués RL, Günther Pomorski T. 2023. Functional Analysis of the P-Type ATPases Apt2-4 from Cryptococcus neoformans by Heterologous Expression in Saccharomyces cerevisiae. J Fungi (Basel) 9.

9. Huang W, Liao G, Baker GM, Wang Y, Lau R, Paderu P, Perlin DS, Xue C. 2016. Lipid Flippase Subunit Cdc50 Mediates Drug Resistance and Virulence in *Cryptococcus neoformans*. mBio 7.

10. Hu G, Caza M, Bakkeren E, Kretschmer M, Bairwa G, Reiner E, Kronstad J. 2017. A P4-ATPase subunit of the Cdc50 family plays a role in iron acquisition and virulence in *Cryptococcus neoformans*. Cell Microbiol 19.

11. Brown HE, Ost KS, Esher SK, Pianalto KM, Saelens JW, Guan Z, Andrew Alspaugh J. 2018. Identifying a novel connection between the fungal plasma membrane and pH-sensing. Mol Microbiol 109:474–493.

12. Chen KZ, Wang LL, Liu JY, Zhao JT, Huang SJ, Xiang MJ. 2023. P4-ATPase subunit Cdc50 plays a role in yeast budding and cell wall integrity in *Candida glabrata*. BMC Microbiol 23:99.

13. Xu D, Zhang X, Zhang B, Zeng X, Mao H, Xu H, Jiang L, Li F. 2019. The lipid flippase subunit Cdc50 is required for antifungal drug resistance, endocytosis, hyphal development and virulence in *Candida albicans*. FEMS Yeast Res 19.

14. Konarzewska P, Wang Y, Han GS, Goh KJ, Gao YG, Carman GM, Xue C. 2019. Phosphatidylserine synthesis is essential for viability of the human fungal pathogen *Cryptococcus neoformans*. J Biol Chem 294:2329–2339.

15. Birge RB, Boeltz S, Kumar S, Carlson J, Wanderley J, Calianese D, Barcinski M, Brekken RA, Huang X, Hutchins JT, Freimark B, Empig C, Mercer J, Schroit AJ, Schett G, Herrmann M. 2016. Phosphatidylserine is a global immunosuppressive signal in efferocytosis, infectious disease, and cancer. Cell Death & Differentiation 23:962–978.

16. Clark GJ, Ju X, Tate C, Hart DN. 2009. The CD300 family of molecules are evolutionarily significant regulators of leukocyte functions. Trends Immunol 30:209–17.

17. Borrego F. 2013. The CD300 molecules: an emerging family of regulators of the immune system. Blood 121:1951–60.

18. Voss OH, Tian L, Murakami Y, Coligan JE, Krzewski K. 2015. Emerging role of CD300 receptors in regulating myeloid cell efferocytosis. Mol Cell Oncol 2:e964625.

19. Rodrigues ML, Janbon G, O’Connell RJ, Chu TT, May RC, Jin H, Reis FCG, Alves LR, Puccia R, Fill TP, Rizzo J, Zamith-Miranda D, Miranda K, Gonçalves T, Ene IV, Kabani M, Anderson M, Gow NAR, Andes DR, Casadevall A, Nosanchuk JD, Nimrichter L. 2025. Characterizing extracellular vesicles of human fungal pathogens. Nat Microbiol 10:825–835.

20. Brandt P, Singha R, Ene Iuliana V. 2024. Hidden allies: how extracellular vesicles drive biofilm formation, stress adaptation, and host–immune interactions in human fungal pathogens. mBio 15:e03045–23.

21. Zhang L, Zhang K, Li H, Coelho C, de Souza Gonçalves D, Fu Man S, Li X, Nakayasu Ernesto S, Kim Y-M, Liao W, Pan W, Casadevall A. 2021. *Cryptococcus neoformans*-Infected Macrophages Release Proinflammatory Extracellular Vesicles: Insight into Their Components by Multi-omics. mBio 12:10.1128/mbio.00279-21.

22. Rizzo J, Wong SSW, Gazi AD, Moyrand F, Chaze T, Commere PH, Novault S, Matondo M, Péhau-Arnaudet G, Reis FCG, Vos M, Alves LR, May RC, Nimrichter L, Rodrigues ML, Aimanianda V, Janbon G. 2021. *Cryptococcus* extracellular vesicles properties and their use as vaccine platforms. J Extracell Vesicles 10:e12129.

23. Rizzo J, Trottier A, Moyrand F, Coppée J-Y, Maufrais C, Zimbres Ana Claudia G, Dang Thi Tuong V, Alanio A, Desnos-Ollivier M, Mouyna I, Péhau-Arnaude G, Commere P-H, Novault S, Ene Iuliana V, Nimrichter L, Rodrigues Marcio L, Janbon G. 2023. Coregulation of extracellular vesicle production and fluconazole susceptibility in Cryptococcus neoformans. mBio 14:e00870–23.

24. Bitencourt Tamires A, Hatanaka O, Pessoni Andre M, Freitas Mateus S, Trentin G, Santos P, Rossi A, Martinez-Rossi Nilce M, Alves Lysangela L, Casadevall A, Rodrigues Marcio L, Almeida F. 2022. Fungal Extracellular Vesicles Are Involved in Intraspecies Intracellular Communication. mBio 13:e03272–21.

25. Hankins HM, Sere YY, Diab NS, Menon AK, Graham TR. 2015. Phosphatidylserine translocation at the yeast trans-Golgi network regulates protein sorting into exocytic vesicles. Mol Biol Cell 26:4674–85.

26. Rizzo J, Stanchev LD, da Silva VKA, Nimrichter L, Pomorski TG, Rodrigues ML. 2019. Role of lipid transporters in fungal physiology and pathogenicity. Comput Struct Biotechnol J 17:1278–1289.

27. Andersen JP, Vestergaard AL, Mikkelsen SA, Mogensen LS, Chalat M, Molday RS. 2016. P4-ATPases as Phospholipid Flippases-Structure, Function, and Enigmas. Front Physiol 7:275.

28. Colombo M, Raposo G, Théry C. 2014. Biogenesis, secretion, and intercellular interactions of exosomes and other extracellular vesicles. Annu Rev Cell Dev Biol 30:255–89.

29. Szule JA, Fuller NL, Rand RP. 2002. The effects of acyl chain length and saturation of diacylglycerols and phosphatidylcholines on membrane monolayer curvature. Biophys J 83:977–84.

30. Nimrichter L, de Souza MM, Del Poeta M, Nosanchuk JD, Joffe L, Tavares PdM, Rodrigues ML. 2016. Extracellular Vesicle-Associated Transitory Cell Wall Components and Their Impact on the Interaction of Fungi with Host Cells. Frontiers in Microbiology Volume 7–2016.

31. Joffe LS, Nimrichter L, Rodrigues ML, Poeta MD. 2016. Potential Roles of Fungal Extracellular Vesicles during Infection. mSphere 1:10.1128/msphere.00099-16.

32. Liu J, Hu X. 2023. Fungal extracellular vesicle-mediated regulation: from virulence factor to clinical application. Frontiers in Microbiology Volume 14 - 2023.

33. Oliveira DL, Freire-de-Lima CG, Nosanchuk JD, Casadevall A, Rodrigues ML, Nimrichter L. 2010. Extracellular vesicles from *Cryptococcus neoformans* modulate macrophage functions. Infect Immun 78:1601–9.

34. Tsou WI, Nguyen KQ, Calarese DA, Garforth SJ, Antes AL, Smirnov SV, Almo SC, Birge RB, Kotenko SV. 2014. Receptor tyrosine kinases, TYRO3, AXL, and MER, demonstrate distinct patterns and complex regulation of ligand-induced activation. J Biol Chem 289:25750–63.

35. Wallet MA, Sen P, Flores RR, Wang Y, Yi Z, Huang Y, Mathews CE, Earp HS, Matsushima G, Wang B. 2008. MerTK is required for apoptotic cell–induced T cell tolerance. The Journal of experimental medicine 205:219–232.

36. Cai B, Thorp EB, Doran AC, Subramanian M, Sansbury BE, Lin C-S, Spite M, Fredman G, Tabas I. 2016. MerTK cleavage limits proresolving mediator biosynthesis and exacerbates tissue inflammation. Proceedings of the National Academy of Sciences 113:6526–6531.

37. Honorato L, Bonilla JJA, Piffer AC, Nimrichter L. 2021. Fungal Extracellular Vesicles as a Potential Strategy for Vaccine Development. Curr Top Microbiol Immunol 432:121–138.

38. Reis FCG, Costa JH, Honorato L, Nimrichter L, Fill TP, Rodrigues ML. 2021. Small Molecule Analysis of Extracellular Vesicles Produced by *Cryptococcus gattii*: Identification of a Tripeptide Controlling Cryptococcal Infection in an Invertebrate Host Model. Front Immunol 12:654574.

39. Piraine REA, Oliveira HT, Santos PW, Froldi JL, Oliveira BTM, Rezende CP, Trentin GES, Nogueira LFB, Colombo AL, Casadevall A, Rodrigues ML, Almeida F. 2025. Cross-Species Communication via Fungal Extracellular Vesicles. bioRxiv doi:10.1101/2025.02.03.636213:2025.02.03.636213.

40. Wolf JM, Espadas J, Luque-Garcia J, Reynolds T, Casadevall A. 2015. Lipid Biosynthetic Genes Affect *Candida albicans* Extracellular Vesicle Morphology, Cargo, and Immunostimulatory Properties. Eukaryot Cell 14:745–54.

41. Rizzo J, Oliveira Débora L, Joffe Luna S, Hu G, Gazos-Lopes F, Fonseca Fernanda L, Almeida Igor C, Frases S, Kronstad James W, Rodrigues Marcio L. 2014. Role of the Apt1 Protein in Polysaccharide Secretion by *Cryptococcus neoformans*. Eukaryotic Cell 13:715–726.

42. Rizzo J, Colombo AC, Zamith-Miranda D, Silva VKA, Allegood JC, Casadevall A, Del Poeta M, Nosanchuk JD, Kronstad JW, Rodrigues ML. 2018. The putative flippase Apt1 is required for intracellular membrane architecture and biosynthesis of polysaccharide and lipids in *Cryptococcus neoformans*. Biochim Biophys Acta Mol Cell Res 1865:532–541.

43. Frallicciardi J, Melcr J, Siginou P, Marrink SJ, Poolman B. 2022. Membrane thickness, lipid phase and sterol type are determining factors in the permeability of membranes to small solutes. Nature Communications 13:1605.

44. Varsanyi C, Birge RB. 2024. Mertk signaling and immune regulation in T cells. J Leukoc Biol 117.

45. Davra V, Kumar S, Geng K, Calianese D, Mehta D, Gadiyar V, Kasikara C, Lahey KC, Chang YJ, Wichroski M, Gao C, De Lorenzo MS, Kotenko SV, Bergsbaken T, Mishra PK, Gause WC, Quigley M, Spires TE, Birge RB. 2021. Axl and Mertk Receptors Cooperate to Promote Breast Cancer Progression by Combined Oncogenic Signaling and Evasion of Host Antitumor Immunity. Cancer Res 81:698–712.

46. Voss OH, Moin I, Gaytan H, Sadik M, Ullah S, Rahman MS. 2025. Phosphatidylserine-binding receptor, CD300f, on macrophages mediates host invasion of pathogenic and non-pathogenic rickettsiae. Infect Immun 93:e0005925.

47. Xu P, Baldridge RD, Chi RJ, Burd CG, Graham TR. 2013. Phosphatidylserine flipping enhances membrane curvature and negative charge required for vesicular transport. J Cell Biol 202:875–86.

48. Morita S-y, Ikeda Y. 2022. Regulation of membrane phospholipid biosynthesis in mammalian cells. Biochemical Pharmacology 206:115296.

49. Chayakulkeeree M, Rude TH, Toffaletti DL, Perfect JR. 2007. Fatty acid synthesis is essential for survival of *Cryptococcus neoformans* and a potential fungicidal target. Antimicrob Agents Chemother 51:3537–45.

50. Leon-Rodriguez CMD, Fu MS, Çorbali MO, Cordero RJB, Casadevall A. 2018. The Capsule of *Cryptococcus neoformans* Modulates Phagosomal pH through Its Acid-Base Properties. mSphere 3:10.1128/msphere.00437-18.

51. DeLeon-Rodriguez CM, Casadevall A. 2016. *Cryptococcus neoformans:* Tripping on Acid in the Phagolysosome. Frontiers in Microbiology Volume 7 - 2016.

52. De Leon-Rodriguez CM, Fu MS, Çorbali MO, Cordero RJB, Casadevall A. 2018. The Capsule of Cryptococcus neoformans Modulates Phagosomal pH through Its Acid-Base Properties. mSphere 3.

53. Tucker SC, Casadevall A. 2002. Replication of *Cryptococcus neoformans* in macrophages is accompanied by phagosomal permeabilization and accumulation of vesicles containing polysaccharide in the cytoplasm. Proc Natl Acad Sci U S A 99:3165–70.

54. Dragotakes Q, Jacobs E, Ramirez LS, Yoon OI, Perez-Stable C, Eden H, Pagnotta J, Vij R, Bergman A, D’Alessio F, Casadevall A. 2022. Bet-hedging antimicrobial strategies in macrophage phagosome acidification drive the dynamics of *Cryptococcus neoformans* intracellular escape mechanisms. PLoS Pathog 18:e1010697.

55. Artavanis-Tsakonas K, Love JC, Ploegh HL, Vyas JM. 2006. Recruitment of CD63 to *Cryptococcus neoformans* phagosomes requires acidification. Proceedings of the National Academy of Sciences 103:15945–15950.

56. Smith LM, Dixon EF, May RC. 2015. The fungal pathogen Cryptococcus neoformans manipulates macrophage phagosome maturation. Cellular Microbiology 17:702–713.

57. Nicola AM, Robertson EJ, Albuquerque P, Derengowski LdS, Casadevall A. 2011. Nonlytic Exocytosis of Cryptococcus neoformans from Macrophages Occurs In Vivo and Is Influenced by Phagosomal pH. mBio 2:10.1128/mbio.00167-11.

58. Stukes SA, Cohen HW, Casadevall A. 2014. Temporal kinetics and quantitative analysis of *Cryptococcus neoformans* nonlytic exocytosis. Infect Immun 82:2059–67.

59. Smith LM, Dixon EF, May RC. 2015. The fungal pathogen *Cryptococcus neoformans* manipulates macrophage phagosome maturation. Cell Microbiol 17:702–13.

60. Santiago-Burgos EJ, Stuckey PV, Santiago-Tirado FH. 2022. Real-time visualization of phagosomal pH manipulation by *Cryptococcus neoformans* in an immune signal-dependent way. Frontiers in Cellular and Infection Microbiology Volume 12–2022.

61. Cao C, Wang Y, Husain S, Soteropoulos P, Xue C. 2019. A Mechanosensitive Channel Governs Lipid Flippase-Mediated Echinocandin Resistance in *Cryptococcus neoformans*. mBio 10.

62. Cao C, Xue C. 2020. More than flipping the lid: Cdc50 contributes to echinocandin resistance by regulating calcium homeostasis in *Cryptococcus neoformans*. Microb Cell 7:115–118.

63. Tancer RJ, Pawar S, Wang Y, Ventura CR, Wiedman G, Xue C. 2024. Improved Broad Spectrum Antifungal Drug Synergies with Cryptomycin, a Cdc50-Inspired Antifungal Peptide. ACS Infect Dis 10:3973–3993.

64. Tancer RJ, Wang Y, Pawar S, Xue C, Wiedman GR. 2022. Development of Antifungal Peptides against *Cryptococcus neoformans*; Leveraging Knowledge about the cdc50Δ Mutant Susceptibility for Lead Compound Development. Microbiol Spectr 10:e0043922.

65. Chen X, Duan HD, Hoy MJ, Koteva K, Spitzer M, Guitor AK, Puumala E, Fiebig AA, Hu G, Yiu B, Chou S, Bian Z, Choi Y, Guo ABY, Wang W, Sun S, Robbins N, Averette AF, Cook MA, Truant R, MacNeil LT, Brown ED, Kronstad JW, Coombes BK, Cowen LE, Heitman J, Li H, Wright GD. 2025. Butyrolactol A enhances caspofungin efficacy via flippase inhibition in drug-resistant fungi. Cell doi:10.1016/j.cell.2025.11.036.

66. Reis FCG, Borges BS, Jozefowicz LJ, Sena BAG, Garcia AWA, Medeiros LC, Martins ST, Honorato L, Schrank A, Vainstein MH, Kmetzsch L, Nimrichter L, Alves LR, Staats CC, Rodrigues ML. 2019. A Novel Protocol for the Isolation of Fungal Extracellular Vesicles Reveals the Participation of a Putative Scramblase in Polysaccharide Export and Capsule Construction in *Cryptococcus gattii*. mSphere 4.

67. Reis Flavia CG, Gimenez B, Jozefowicz Luísa J, Castelli Rafael F, Martins Sharon T, Alves Lysangela R, de Oliveira Haroldo C, Rodrigues Marcio L. 2021. Analysis of Cryptococcal Extracellular Vesicles: Experimental Approaches for Studying Their Diversity Among Multiple Isolates, Kinetics of Production, Methods of Separation, and Detection in Cultures of Titan Cells. Microbiology Spectrum 9:10.1128/spectrum.00125-21.

68. Nicola AM, Frases S, Casadevall A. 2009. Lipophilic dye staining of *Cryptococcus neoformans* extracellular vesicles and capsule. Eukaryot Cell 8:1373–80.

